# A high throughput assay provides novel insight into type VII secretion in *Staphylococcus aureus*

**DOI:** 10.1101/2023.06.03.543475

**Authors:** Yaping Yang, Aaron A. Scott, Holger Kneuper, Felicity Alcock, Tracy Palmer

## Abstract

Successful colonisation by the opportunistic pathogen *Staphylococcus aureus* depends on its ability to interact with other microorganisms. *S. aureus* strains harbour a T7b-subtype type VII secretion system (T7SSb), a protein secretion system found in a wide variety of Bacillota which functions in bacterial antagonism and virulence. Assessment of T7SSb activity in *S. aureus* has been hampered by low secretion activity under laboratory conditions, and the lack of a sensitive assay to measure secretion. Here we have utilised NanoLuc Binary Technology to develop a simple assay to monitor protein secretion via detection of bioluminescence. Fusion of the 11 amino acid NanoLuc fragment to the conserved substrate EsxA permits its extracellular detection upon supplementation with the large NanoLuc fragment and luciferase substrate. Following miniaturisation of the assay to 384 well format, we use high-throughput analysis to demonstrate that T7SSb-dependent protein secretion differs across strains and growth temperature. We further show that the same assay can be used to monitor secretion of the surface-associated toxin substrate TspA. Using this approach we identify three conserved accessory proteins required to mediate TspA secretion. Co-purification experiments confirm that all three proteins form a complex with TspA.

## Background

The opportunistic pathogen *Staphylococcus aureus* is a major cause of both nosocomial and community infections. It is a frequent coloniser of animals and humans and infections can result in a range of life-threatening diseases such as pneumonia, endocarditis and sepsis. It is notorious for developing resistance to frontline antimicrobials, with methicillin resistance now widespread and vancomycin resistance also increasing [1, 2]. Like many Gram-positive bacteria, *S. aureus* carries a specialised protein secretion machinery called the type VII secretion system (T7SS), which it utilises to secrete antibacterial toxins targeting competitor bacteria (reviewed in [3]). T7SS-mediated antagonism is likely to be important for colonisation [4], and the *S. aureus* T7SS has been shown to be required for virulence in mouse infection models [5–9]. There are several distinct T7SS subtypes, of which the T7SSa and T7SSb are the best characterised [10]. The T7SSb is widely distributed among Bacillota, and has been studied to date in species of *Streptococcus*, *Staphylococcus*, *Bacillus* and *Enterococcus* [8, 11–13]. However, in many cases functional and mechanistic studies have been hindered by low secretion activity in laboratory conditions.

The *S. aureus* T7SSb is encoded on the core genome at the *ess* locus, and four variant locus types have been identified, termed *essC-1* – *essC-4*. The genes at the 5’ end of the *ess* locus are highly conserved and encode core components of the secretion machinery, while the 3’ end codes for different complements of variant-specific substrate and accessory genes [4, 14]. The primary component of the T7SSb transmembrane channel is the ATPase EssC, inferred from its homology to the EccC component of the mycobacterial T7SSa channel whose structure has been solved by cryo-electron microscopy [15, 16]. EssC has a conserved N-terminus, two transmembrane domains which likely form the secretion channel, and four C-terminal ATPase domains, the latter two of which are variant-specific and provide substrate specificity [17]. Three additional membrane proteins, EsaA, EssA and EssB, and two small globular proteins, EsxA and EsaB, are further essential components of the T7SSb secretion system [6, 8, 18].

In addition to being a core component of the T7SSb secretion system, EsxA is itself a T7SS secretion substrate. EsxA secretion is essential for the export of other T7SSb substrates, and it appears to be co-secreted with at least some of them [8, 19]. EsxA is a member of the WXG100 protein family, which are helical hairpin proteins that often, but not always, harbour a central W-x-G sequence motif [20, 21]. In addition to EsxA, other WXG100-like proteins, with more specialised functions, are also found in T7SSb^+^ strains. These are usually encoded at genetic loci with genes for larger T7SSb-secreted toxins. T7SSb toxins contain an LXG domain, which can be located either at the protein N-terminus (for example TelB, EsaD and TspA) or C-terminus (e.g. TslA) [6, 11, 18, 22–25]. Prior to secretion, the co-encoded WXG100-like proteins, which have been termed Laps (for <u>L</u>XG-<u>a</u>ssociated α-helical <u>p</u>roteins) bind to the LXG domain of the cognate toxin partner. This interaction generates a secretion-competent rod-shaped complex, and conserved sequence motifs on both the LXG domain and one of the Laps constitute a T7SSb-targeting signal [23, 24, 26, 27]. In some instances, a globular protein is also encoded at the toxin genetic locus, which also facilitates toxin secretion [24, 27].

TspA is a membrane-depolarising toxin with antibacterial activity that is encoded by all *S. aureus* strains analysed to date. Following secretion, TspA associates with the cell surface and can only be released experimentally by digestion of the cell wall [22]. Unlike other characterised LXG toxins, no WXG100-like partner proteins are encoded at the *tspA* locus, and its secretion requirements are unclear. Assessing secretion by the *S. aureus* T7SS is hampered by low secretion activity in laboratory growth media. Moreover, the lack of cleavable signal peptides on T7SSb substrates means there is no size difference between the exported and cytosolic forms making cellular lysis a confounding issue, particularly where cell wall digestion is also required. To circumvent these difficulties, we have developed a novel secretion assay that makes use of NanoLuc binary technology [28]. We show that this is a robust reporter system that can be used to monitor both secreted and cell-surface localised T7SSb substrates and which is amenable to high-throughput analysis. Using this assay we identify three genes that are required for efficient export of TspA, and demonstrate that the encoded proteins directly interact with the toxin.

## Methods

### Bacterial strains, plasmids and growth conditions

*E. coli* was cultured in LB medium (Melford) and *S. aureus* was cultured in tryptic soy broth (TSB; Oxoid) with chloramphenicol (10 μg/ml) where required. *E. coli* strains DH5α (*Δ(argF-lac)169, φ80dlacZ58(M15), ΔphoA8, glnX44(AS), deoR481, rfbC1, gyrA96(NalR), recA1, endA1, thiE1 and* hsdR17) and JM110 *rpsL thr leu thi lacY galK galT ara tonA tsx dam dcm glnV44 Δ(lac-proAB)* e14-[F’ *traD36 proAB^+^ lacI^q^ lacZ*ΔM15] *hsdR17*(rK^−^mK*^+^*) were used for cloning and preparation of plasmids for electroporation, respectively, and M15 harbouring pREP4 (*F-, lac, ara, gal, mtl* [*KanR, lacI*]) was used for protein overproduction. *S. aureus* strains and plasmids used are listed in Tables 1 and 2, respectively. Chromosomal deletion of *essC* was accomplished by allelic exchange using plasmid pIMAY [29] carrying the *essC* flanking regions. Insertion of the pep86 tag on the chromosome of COL and COLΔ*essC* used an updated system with plasmid pIMAY-Z [30]. pIMAY plasmids were created using standard restriction cloning techniques, and pIMAY-Z-esxApep86 was created using HiFi assembly (NEB). Plasmids pRab11-esxApep86 and pRab11-tspA_1-328_-pep86_2603_2602_2601 for NanoLuc secretion assays, and plasmid pQE70-_Tstrep_SACOL2603_myc_-2602-2601_HA_-_his_TspA for protein purification were created using HiFi assembly (NEB, E0554S). *SACOL2603*, *SACOL2602* and *SACOL2601* were individually deleted from pRab11-tspA_1-328_-pep86_2603_2602_2601 using the Q5 site-directed mutagenesis kit (NEB). Oligonucleotides are listed in Table S1.

**Table 1.** *S. aureus* strains used in this study.

| Strain | Description | Reference |
| --- | --- | --- |
| COL | MRSA, *agr*, *essC*-1 variant strain | [45, 46] |
| COLΔ*essC* | COL with markerless deletion of *essC* | [22] |
| RN6390 | NCTC8325 derivative. *rbsU*, *tcaR*, cured of ϕ11, ϕ12, ϕ13. *essC*-1 variant strain | [47, 48] |
| RN6390Δ*essC* | RN6390 with markerless deletion of *essC* | [8] |
| JP5347 | Human isolate, *essC*-1 variant strain | This work |
| JP5347Δ*essC* | JP5347 with markerless deletion of *essC* | This work |
| MRSA252 | Nosocomial HA-MRSA isolate, representative of Epidemic MRSA-16. *essC*-2 variant strain | [49] |
| MRSA252Δ*essC* | MRSA252 with markerless deletion of *essC* | This work |
| 10.1252.X | Livestock-associated ST398 isolate. *essC*-3 variant strain | [33] |
| 10.1252.XΔ*essC* | 10.1252.X with markerless deletion of *essC* | This work |
| EMRSA15 | EMRSA-15 clonal complex 22 human isolate. *essC*-4 variant strain. | [50] |
| EMRSA15Δ*essC* | EMRSA15 with markerless deletion of *essC* | This work |
| COL<sub>*esxA-pep86*</sub> | Pep86 coding sequence (VSGWRLFKKIS) fused to the 3' end of *esxA* in COL | This work |
| COLΔ*essC*<sub>*esxA-pep86*</sub> | Pep86 coding sequence (VSGWRLFKKIS) fused to the 3' end of *esxA* in COLΔ*essC* | This work |

**Table 2.**
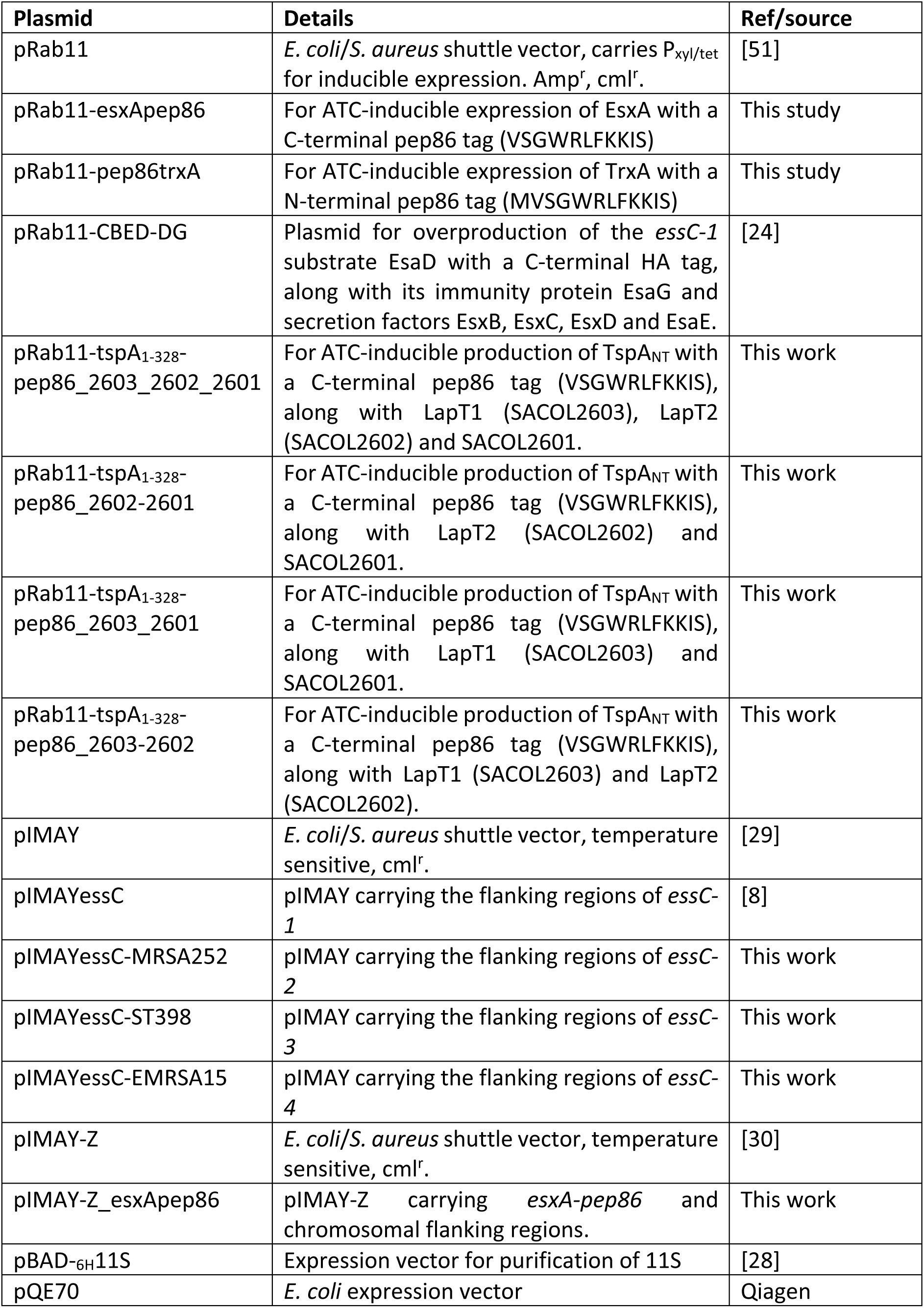
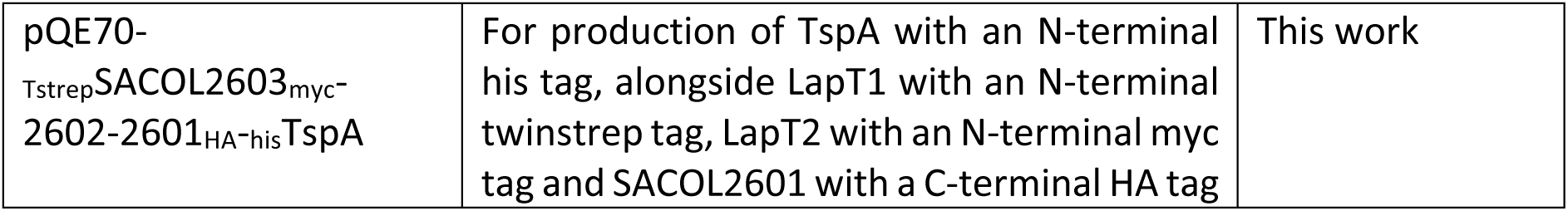
Plasmids used in this study.

### NanoLuc secretion assays

Furimazine was purchased from Promega (N1610) and used according to the manufacturer’s instructions. The NanoBit large subunit 11S was purified from *E. coli* BL21(DE3) [pBAD-_6H_11S] using the published protocol [28] and was used at a final concentration of 5 μM. For assay of 5 ml cultures, *S. aureus* strain COL were cultured as described above. Once cultures had reached a density of 0.5 (OD_600_), anhydrotetracycline (ATC; 250ng/ml) was added to induce expression of EsxA-pep86. Cells were harvested at a density of 2 (OD_600_) by centrifugation at 16,000 x g, 20°C. Cell pellets were resuspended in TBS + 50μg/ml lysostaphin (LSPN, Ambi) and incubated for 10 minutes at 37°C. 100μl of each sample was supplemented with 5µM 11S and 2µl furimazine (1:100 dilution of stock solution) in a 96-well plate (Greiner, REF655073) with an approximate final well volume of 105μl. Luminescence at 460 nm was read 3 minutes after incubation of the plate at room temperature, with a FLUOstar Omega using a gain value of 3,000. Samples from the same cultures were prepared for immunoblot and analysed with antibodies as previously described [24]. For assays in 384-well plates, overnight *S. aureus* cultures were subcultured in fresh TSB at a density (OD_600_) of 0.001 for assay of EsxA-pep86 secretion or 0.01 for assay of TspA_NT_-pep86 secretion and cultured at 37 °C/34 °C/30 °C with shaking at 200rpm in a 384-well plate (Greiner, REF781098), with a well volume of 50μl. In general, for each assay one row of the 384 well plate was used (16 wells, i.e. 16 technical replicates). This was repeated in triplicate with three separate starter cultures (i.e. three biological replicates). The exceptions to this are in Fig. 2 D and E, where in Fig 2D, there is one biological replicate but 64 technical replicates and Fig 2E where there are three biological repeats each of 64 technical replicates. For COL strains grown at 37 °C, 250 ng/ml ATC was added after 320 min and 11S and furimazine were added after 380 min. For all other strains 500 ng/ml ATC was added after 180 minutes (37°C cultures), 220 minutes (34°C cultures), or 240 minutes (30°C cultures). 11S and furimazine were added at 220 minutes (37°C cultures), 280 minutes (34°C cultures) or 300 minutes (30°C cultures). OD_600_ and luminescence readings were taken every 10 minutes. To measure total luminescence (i.e. combined extracellular and cytoplasmic levels), 50μg/ml lysostaphin in TBS was added to each well and incubated for 10 minutes at 37°C. luminescence readings were taken after supplementation of 11S and furimazine.

**Figure 1.**
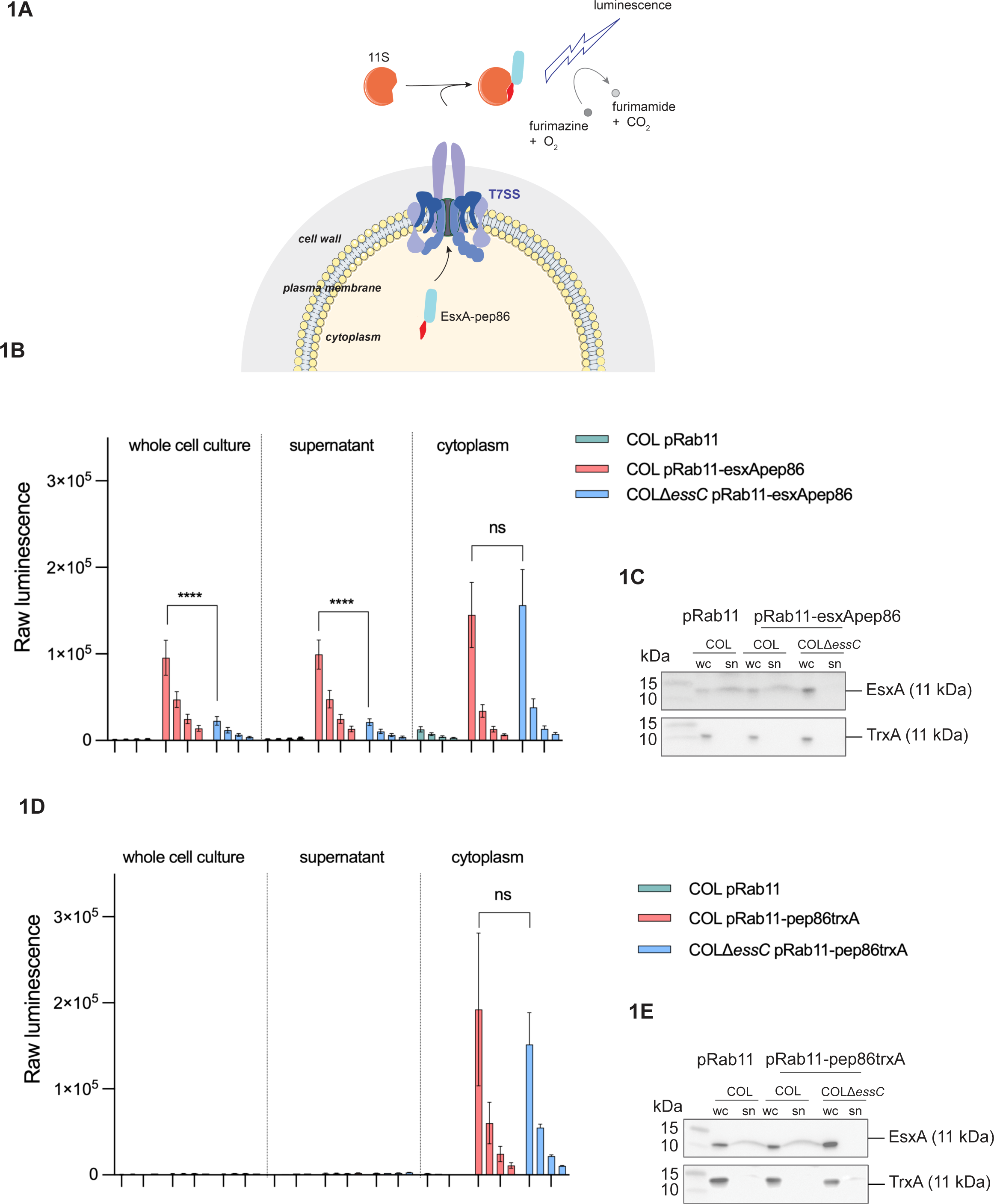
The NanoBit T7-secretion assay. A. Schematic representation of the NanoBit secretion assay. The T7SSb substrate EsxA with a C-terminal pep86 tag is produced in the *S. aureus* cytoplasm, and 11S and furimazine (the NanoLuc large subunit and substrate) are provided outside the cell. Secretion of EsxA-pep86 via the T7SS results in functional complementation of 11S, which luminesces in the presence of furimazine and O_2_. Note that the cartoon is not an accurate representation of the stoichiometry and architecture of the T7SSb, which are unknown. B, D. COL wild-type or Δ*essC* strains carrying the indicated plasmids were grown in 5 ml cultures. At OD_600_ = 0.5 ATC was added to induce gene expression from pRab11. When cells reached OD_600_ = 2, samples corresponding to the whole cell culture, the clarified culture medium (‘supernatant’) and lysed cells (‘cytoplasm’) were prepared. Serial two-fold dilutions of whole cell cultures, clarified supernatant and cytoplasmic fractions were prepared, and 11S and furimazine were added to each. The data presented correspond to the mean luminescence readings of five biological replicates and error bars represent the SEM. Two-way ANOVA was performed to determine statistical significance (ns p>0.05; **** p<0.0001). C, E. Lysed whole cell (wc) and clarified culture supernatant (sn) fractions from (B) and (D) were analysed by immunoblot with antibodies against EsxA (top) or TrxA (bottom). Supernatant samples were concentrated by TCA precipitation prior to analysis, and the amount analysed corresponds to 6 x more culture volume than for lysed cell samples.

**Figure 2.**
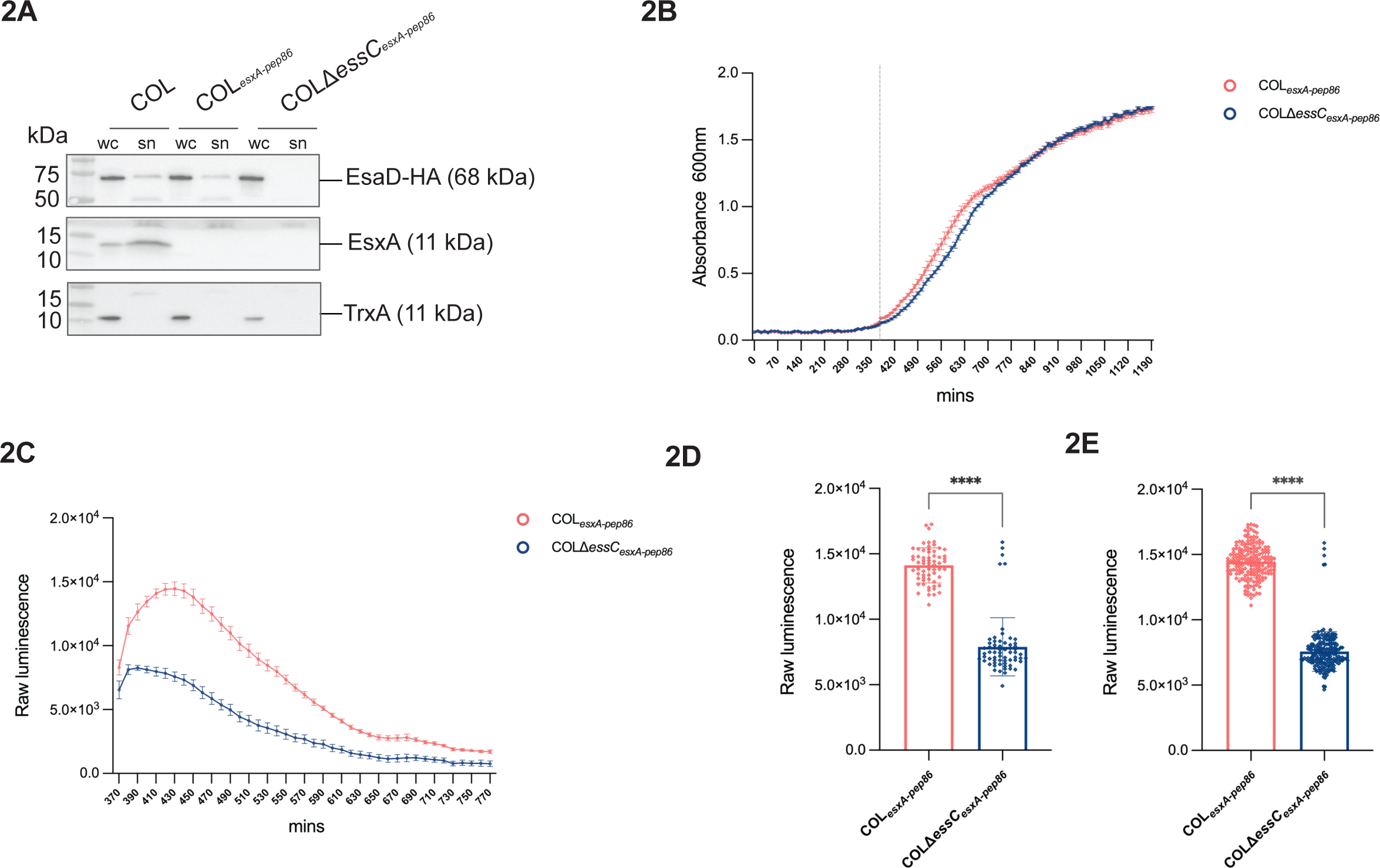
**High throughput secretion analysis of strains producing chromosomally-encoded EsxA-pep86.** A. The indicated strains were grown in 5 ml cultures and ATC was added at OD_600_ = 0.5 to induce gene expression from pRab11. ‘pEsaD-HA’ denotes plasmid pRab11-CBED-DEG, carrying an HA-tagged copy of the T7 substrate EsaD, along with its accessory secretion factors [24]. At OD_600_ = 2, lysed whole cell (wc) and clarified supernatant (sn) fractions were prepared and analysed by immunoblot with antibodies against the HA tag (to detect EsaD-HA), EsxA or TrxA. Supernatant samples were concentrated by TCA precipitation prior to analysis, and the amount analysed corresponds to 6 x more culture volume than for lysed cell samples. B. Growth curve for cultures of COL wild-type and Δ*essC* strains, each producing chromosomally encoded EsxA with a C-terminal pep86 tag from the native locus. 192 replicate 50 μl cultures (64 technical replicates x 3 biological replicates) were grown in 384-well plates at 37°C. Furimazine and 11S were added at 360 min (dotted line). Mean OD_600_ values are plotted and error bars correspond to the SEM (n=3). C. Luminescence readings at ten minute intervals for the cultures in (B) following initiation of luminescence with furimazine and 11S at 360 minutes. Error bars correspond to the SEM (n=3). D. A single overnight culture of each COL*_esxA-pep86_* and COLΔ*essC_esxA-pep86_* was diluted to an OD_600_ of 0.1 and used to inoculate 64 individual wells of a 384 well plate. After 360 minutes growth, furimazine and 11S were added and a single luminescence reading from each well was taken at 430 minutes. E. As for (D) except that three overnight cultures of each COL*_esxA-pep86_* and COL Δ*essC_esxA-pep86_* were used, giving 192 individual data points for each strain. Statistical significance in (D) and (E) were assessed with the Mann-Whitney test, with **** indicating p< 0.0001 for each case.

**Figure 3.**
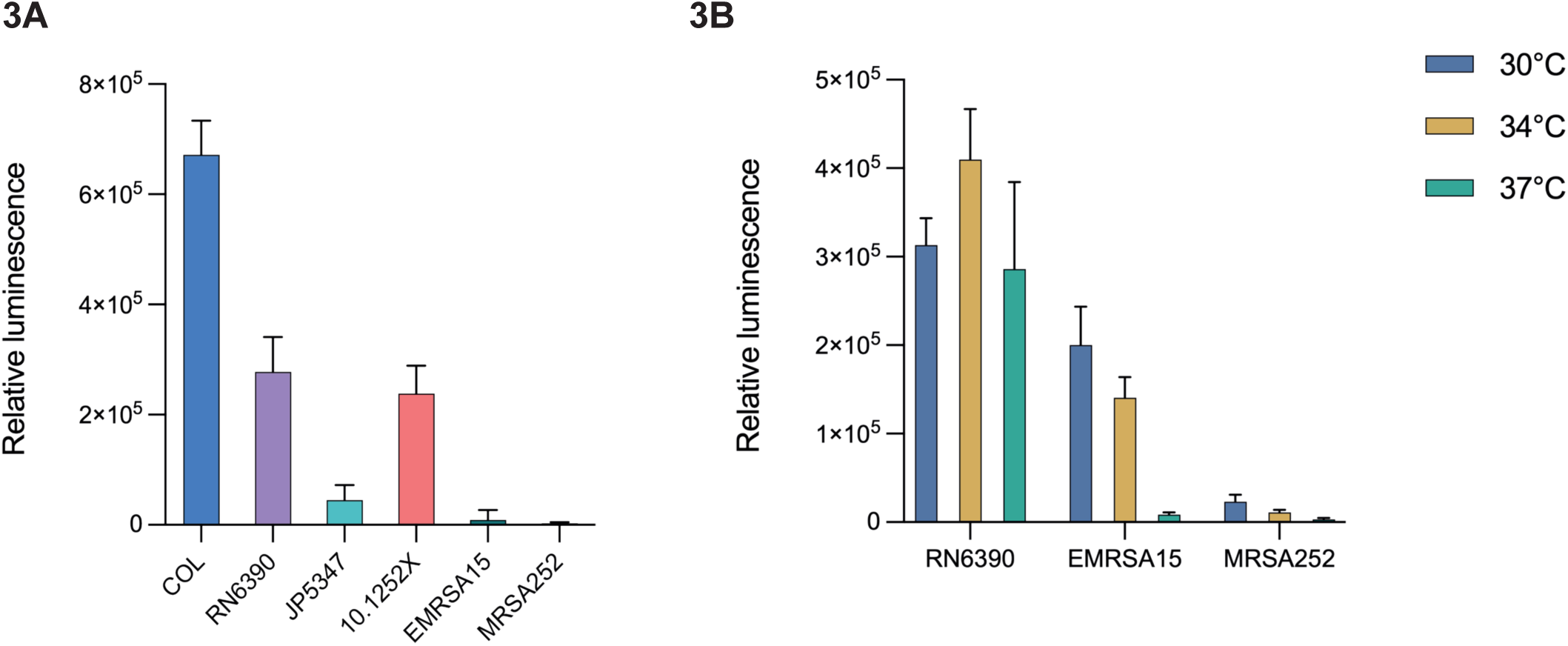
EsxA secretion in different S. aureus strains and at different temperatures. A. Wild-type and Δ*essC* variants of the indicated strains were transformed with pRab11-esxApep86 and cultured in 384-well plates at 37°C. Expression of *esxA-pep86* was induced by addition of ATC at 180 minutes, and luminescence was initiated by addition of furimazine and 11S at 220 minutes. Luminescence readings taken at ten minute intervals (shown in Fig. S1B) were divided by OD_600_ readings to give the relative luminescence. The values for Δ*essC* relative luminescence were subtracted from those of the wild-type relative luminescence to give a relative secretion value for each time point. For each strain pair the time point with the peak value for relative secretion is presented. Data represent the mean of three biological replicates each with 16 technical replicates. Error bars correspond to the SEM (n=3). B. Secretion assays were performed as in (A), but each strain pair was cultured at 30°C, 34°C and 37°C (data for 37°C is reproduced from (A) for the sake of comparison). The timings of ATC induction and luminescence initiation were adjusted for the slower growth rates at 30°C and 34°C (see Fig S2A and methods) and raw luminescence readings are shown in Figure S2B.

### Cell fractionation and western blot

Single colonies of *S. aureus* harbouring the appropriate pRab11-based plasmid were cultured in TSB medium with chloramphenicol. The following day, 600 µl of overnight culture was sub-cultured into 10 ml of fresh TSB. When the culture reached OD_600nm_ = 0.5-0.6, 500 ng/ml of ATC was added to induce plasmid-encoded gene expression. Cells were harvested at OD_600_ = 2. A 1.8 ml aliquot of the culture supernatant was retained, filtered through a 0.22µM sterile filter and the filtrate supplemented with 50 µg/ml sodium deoxycholate (Merck, 89904) and 10% TCA (Cambridge Bioscience, 700016-10 ml-CAY). Proteins were precipitated on ice overnight and subsequently pelleted by centrifugation. After washing with 80% ice-cold ethanol, the pellet was resuspended in resuspension buffer (50mM Tris-HCl, pH 8.0, 4% SDS, 10% glycerol) then boiled with 1/3 volume of 4x Laemmli sample buffer (Bio-Rad, 1610747) for 10 mins. This was taken as the supernatant fraction. The harvested cells were resuspended in Tris-buffered saline containing 100 µg/ml lysostaphin and incubated at 37 °C for 30 mins. This was then supplemented with 1/3 volume of 4x Laemmli sample buffer and boiled for 10 min. This was taken as the whole cell fraction.

For Western blot analysis, samples were separated by SDS PAGE, transferred to nitrocellulose membrane using a Trans-Blot (Bio-Rad) with Whatman paper soaked in transfer buffer composed of 25 mM Tris, 192 mM glycine pH 8.3, 10% methanol, and analysed by incubation with one of the following primary antibodies: anti-EsxA [8], anti-TrxA [31], anti-HA tag (Merck, H9658), anti-strep tag (Qiagen, 34850), anti-6*his tag (Thermo Fisher Scientific,11533923), anti-myc tag (Abcam, Ab23), and an HRP-linked secondary antibody. A goat anti-mouse antibody (Bio-Rad, 1706516) was used with primary antibodies against the strep, myc and HA tags. A goat anti-rabbit antibody (Bio-Rad, 1721019) was used with all other primary antibodies. All uncropped western blots are provided in the supplementary materials.

### Protein purification

LB medium containing ampicillin (0.125 mg/ml) was inoculated 1/100 with an overnight preculture of strain M15 strain carrying pQE70-_Tstrep_SACOL2603-_myc_2602-2601_HA_-_his_TspA and grown at 37°C for 2 to 3 h with aeration, until OD_600_ reached ∼0.5. Cultures were then supplemented with 0.5 mM isopropyl β-d-1-thiogalactopyranoside (IPTG), and grown at 18°C overnight, then harvested by centrifugation. Cell pellets were resuspended in binding buffer W (100 mM Tris/HCl, pH 8.0, 150 mM NaCl, and 1 mM EDTA). Cells were lysed by sonication, the resulting cell lysate was clarified by centrifugation at 20,000 g for 30 min, and the supernatant was applied to a 1ml Strep-tactin XT column (IBA, 2-4021-001), washed with bugger W (100 mM Tris-HCl, pH 8.0, 150 mM NaCl, 1 mM EDTA) and the bound proteins were eluted in buffer BXT (IBA, 2-1042-025). Proteins eluted from the Strep-tactin XT column were concentrated using a 50 MWCO Vivaspin centrifugal concentrator (Global Life Sciences Solutions Operations, 28932318) before injection onto a Superdex S200 10/300 GL column (Cytiva) equilibrated in 50 mM HEPES pH 7.5, 150 mM NaCl. Samples were analysed by SDS PAGE using Mini-PROTEAN® TGX™ Precast Gels (Bio-Rad).

### Other methods

Secretion data were analysed and presented using Prism (Graphad). Raw data is available for download from FigShare (https://doi.org/10.6084/m9.figshare.25368598). Gene neighbourhood digrammes were generated using clinker (https://cagecat.bioinformatics.nl; [32]).

## Results

### A NanoLuc assay for measuring Type VII-dependent secretion in Staphylococcus aureus

Previous methods to monitor *S. aureus* T7SSb secretion activity have relied almost exclusively on western blotting. Assessing secretion of endogenous WXG100-family proteins, in particular EsxA, involves isolating and concentrating culture supernatants, usually by precipitation with trichloroacetic acid [6, 8, 19, 33]. Furthermore, detection of natively produced LXG toxins is generally too low to detect by western blotting even from concentrated supernatant samples, and assessing their secretion has required overproduction of epitope-tagged variants from an inducible plasmid [22–25]. These methods depend on the availability of good quality antibodies, are often poorly reproducible, and are difficult to quantify.

To circumvent these issues we utilised NanoLuc binary technology (abbreviated to NanoBit) to develop a new assay to monitor T7 secretion. NanoLuc is an engineered variant of a deep sea shrimp luciferase protein. It is relatively small (19 kDa) and has superior biophysical characteristics compared to other luciferases [34]. NanoBit is a complementation reporter system which has previously been used to investigate protein secretion via the Sec and Tat translocons in *E. coli* [28, 35, 36]. Two separate NanoLuc subunits (11S, a 156 amino acid large subunit and pep86, an 11 amino acid small subunit) combine with high affinity to generate the functional enzyme, which luminesces with high intensity at 460 nm in the presence of the substrate furimazine and molecular oxygen [34].

We first selected EsxA as our model substrate (Fig 1A). EsxA is produced at significantly higher copy than other T7SS components [8] and can be detected by Western blot in culture supernatants of the *essC-1* strain COL relatively reliably, allowing us to correlate our new assay with data from traditional immunoblot assays. We constructed plasmid pRab11-esxApep86 which permits inducible overproduction of EsxA with the NanoBit small subunit, pep86, fused to its C-terminus. This plasmid was transferred into wild-type and Δ*essC* variants of COL, and the wild-type strain was also transformed with empty pRab11 as a further control. Cultures (5ml volume in 50ml falcon tubes) were supplemented with anhydrotetracycline (ATC) and grown at 37°C to mid logarithmic phase (OD_600_ ≅ 2.0). At this point samples of whole cell culture, culture supernatant and lysostaphin-lysed cells were prepared. The NanoBit large subunit, 11S, and the NanoLuc substrate furimazine were added to serial two-fold dilutions of these samples, and luminescence readings were taken (Fig. 1B). Cell lysate and culture supernatant samples from the same cultures were also analysed by immunoblot (Fig. 1C).

Luminescence was detected in all samples except those from cells carrying empty vector, signifying an EsxA-pep86–specific luminescence signal (Fig. 1B). Similar luminescence readings were obtained for both the whole cell culture and the supernatant samples, demonstrating that external pep86 can be measured without centrifugal separation of culture supernatants (Fig. 1B). Deletion of *essC* resulted in a dramatic reduction in the level of extracellular EsxA-pep86 compared to the wild-type strain, while the cytoplasmic level was similar in both strains. This confirms that the extracellular EsxA-pep86-dependent luminescence signal is due to secretion by the T7SS. We assign that residual low level of extracellular EsxA-pep86 in the *essC* mutant strain to cell leakage/lysis.

When fractions of the same cultures were analysed by immunoblot a single band at ∼11 kDa was detected with antibodies against EsxA in all three strains (Fig. 1C), regardless of whether the cells harboured pRab11 or pRab11-esxApep86. Since two of the three strains analysed produce both endogenous chromosomally-encoded EsxA (11 kDa) and plasmid-encoded EsxA-pep86 (12.3 kDa), we infer that this 11 kDa band corresponds to endogenous EsxA, and that the pep86 fusion inhibits detection by the EsxA antibodies (see also Fig. 2C where chromosomal tagging of *esxA* with pep86 inhibits antibody detection).

As an additional control, we constructed a similar fusion of a cytoplasmic protein, TrxA with an N-terminal pep86 tag (pRab11-pep86-TrxA) and assayed this in the same way. Luminescence readings showed that while pep86-TrxA was produced, barely any signal was detected outside the cells (Fig. 1D). Immunoblots confirmed the cytoplasmic localisation of TrxA and showed that EsxA was secreted normally in these strains, although similarly to EsxA-pep86, the pep86-TrxA fusion was not detected by our TrxA antibodies (Fig. 1E). In summary, Figure 1 demonstrates that luminescence readings correlate with T7-dependent secretion of EsxA-pep86, and that extracellular EsxA-pep86 can be measured in unprocessed whole cell cultures.

### High throughput analysis of EsxA secretion

Having established that we can use the NanoBit assay to measure T7SS secretion in *S. aureus*, and that we can directly detect extracellular EsxA-pep86 from growing cultures, we next sought to miniaturise it for high throughput format to improve quantitation and sensitivity. To this end we engineered a C-terminal pep86 fusion onto chromosomally-encoded EsxA, to avoid the need for an inducible plasmid. This ensures the assay is undertaken using a system that is as close to native as possible. To assess whether the chromosomally-encoded EsxA-pep86 fusion was competent to support T7 secretion, we transformed this strain with a plasmid encoding a HA-tagged copy of the T7 LXG toxin substrate EsaD alongside its accessory proteins. Immunoblot data for cell lysate and culture supernatant samples (Fig. 2A) showed that although the EsxA-pep86 fusion was not detected by our antibodies, as we had seen previously with the plasmid-encoded fusion, secretion of EsaD-HA was unaffected by the presence of the pep86 tag on EsxA. We conclude that EsxA-pep86 is functional (Fig. 2A).

We next performed growth curves for 50 μl cultures of COL*_esxApep86_* and an otherwise isogenic *essC* mutant in 384-well plate format. Strains grew slowly under these conditions, with cultures reaching stationary phase approximately 20 hours (1200 minutes) after inoculation at an OD_600_ of 0.001 (Fig. 2B). The *essC* mutant strain grew slightly more slowly in exponential phase but reached the same final OD_600_ as the *essC*^+^ strain. The half-life of the NanoLuc luminescence signal is reported to be approximately 120 minutes (Promega) meaning that continuous monitoring of secreted EsxA-pep86 throughout exponential growth would not be possible. We therefore chose to monitor secretion during early exponential phase, and to this end furimazine and 11S were added to cultures at the beginning of exponential phase, after six hours (360 minutes) growth (Fig. 2B, dotted line) to initiate luminescence. Luminescence readings were then taken at ten minute intervals. The luminescence signal for COL*_esxApep86_* cultures initially rose, peaking 70 minutes after initiation of luminescence, indicating that cells were actively secreting protein during this time (Fig. 2C). The subsequent decline in luminescence intensity is likely due to signal decay, although degradation of pep86 might also contribute. As seen in the previous assay shown in Figure 1, some leakage of EsxA-pep86 occurred in the Δ*essC* strain (Fig. 2C).

To analyse the distribution of secretion activity within a clonal population, a single colony of each of COL*_esxApep86_* and COLΔ*essC_esxApep86_* was used to inoculate overnight cultures, and these were each used to inoculate 64 x 50 μl cultures and assayed as described above. Luminescence readings taken at 430 minutes (OD_600_ = 0.23) were used to generate a scatter plot of individual data points (Fig. 2D). This analysis reveals a moderate distribution of secretion activity in replicate cultures of COL*_esxApep86_*, however the distribution of extracellular EsxA-pep86 levels in the control COLΔ*essC_esxApep86_* strain was quite broad, overlapping the *essC*^+^ strain and highlighting the requirement for a high throughput assay. The same analysis was repeated for three independent cultures of each strain and this did not increase variability (Fig. 2E).

### Variable secretion characteristics of different *S. aureus* strains

It has previously been reported that T7SS activity varies across different *S. aureus* isolates and growth conditions [8, 19, 33]. However direct comparison across different *S. aureus* strains has been hampered by low secretion activity (often below the detection limit) and low reproducibility (our unpublished observations). We therefore used the NanoBit assay to compare secretion across cell cultures of five additional *S. aureus* isolates, two of which (like COL) carry the *essC-1* variant *ess* locus (RN6390 and JP5347), and one representative isolate for each of *essC-2* (10.125.2X), *essC-3* (MRSA252) and *essC-4* (EMRSA15). Each strain, along with its otherwise isogenic Δ*essC* mutant, was transformed with pRab11-esxApep86 and assayed for EsxA-pep86 secretion in 384-well plates. Growth curves showed that each Δ*essC* strain grew similarly to its parental strain (Fig. S1A). The five additional strains tested here all entered exponential phase faster than COL, and so the timing of addition of ATC (to induce EsxA-pep86 production) and of 11S and furimazine was adjusted accordingly (see methods). T7SS-dependent secretion of EsxA-pep86 was observed, but to varying degree, in the three *essC-1* isolates and the *essC-2* strain 10.1252.X. However no T7-secreted EsxA could be detected in the *essC-3* (MRSA252) and *essC-4* (EMRSA-15) strains (Fig. 3A, S1B). To confirm that EsxA-pep86 was being produced in all of the strains we measured total luminescence in each culture following in-well cellular lysis (a combined measure of cytoplasmic and secreted EsxA-pep86 levels (Fig. S1C). Substantive luminescence could be detected even in lysed cultures of MRSA252 and EMRSA-15, indicating that these strains produce but do not secrete EsxA-pep86.

Since no T7SS activity was detected for strains MRSA252 and EMRSA15 grown at 37°C we wondered whether these isolates might exhibit a different temperature dependence for secretion. We therefore repeated the NanoBit assay with these strains, alongside RN6390 as an *essC-1* control, using cell cultures growing at 30°C and 34°C (Fig. 3B and S2). For EMRSA15, EsxA-pep86 was secreted at both of the lower temperatures, while for MRSA252 the lower temperatures made little difference to secretion activity, which was negligible at all temperatures tested. EsxA was secreted similarly at all three temperatures in RN6390 (Fig. 3B and S2). Temperature therefore contributes to the regulation of T7SS activity in *S. aureus* in a strain-specific manner. Of note, EsxA-pep86 leakage was markedly lower at 30°C and 34°C than at 37°C in each of the Δ*essC* strains (Fig. S2B).

### Identification of TspA secretion partners

TspA is a membrane-depolarising LXG toxin found across all *S. aureus essC* variant strains. It is encoded at a locus remote from the T7SS structural genes and precedes a string of non-identical genes coding for DUF443 membrane proteins which provide protection against TspA activity [22] (Fig. 4A). Recent analysis of other T7SSb LXG toxins has revealed a common mechanistic feature for secretion, involving formation of a hetero-oligomeric pre-secretion complex between the LXG domain and helical hairpin partner proteins encoded at the same genetic locus as the toxin [24, 26, 27, 37]. Logically, the TspA LXG domain should also be expected to require similar helical binding partners to generate a secretion-competent structure. However, there are no candidate genes coding for predicted helical proteins in the immediate neighbourhood of *tspA* (*SACOL0643*) (Fig. 4A).

**Figure 4.**
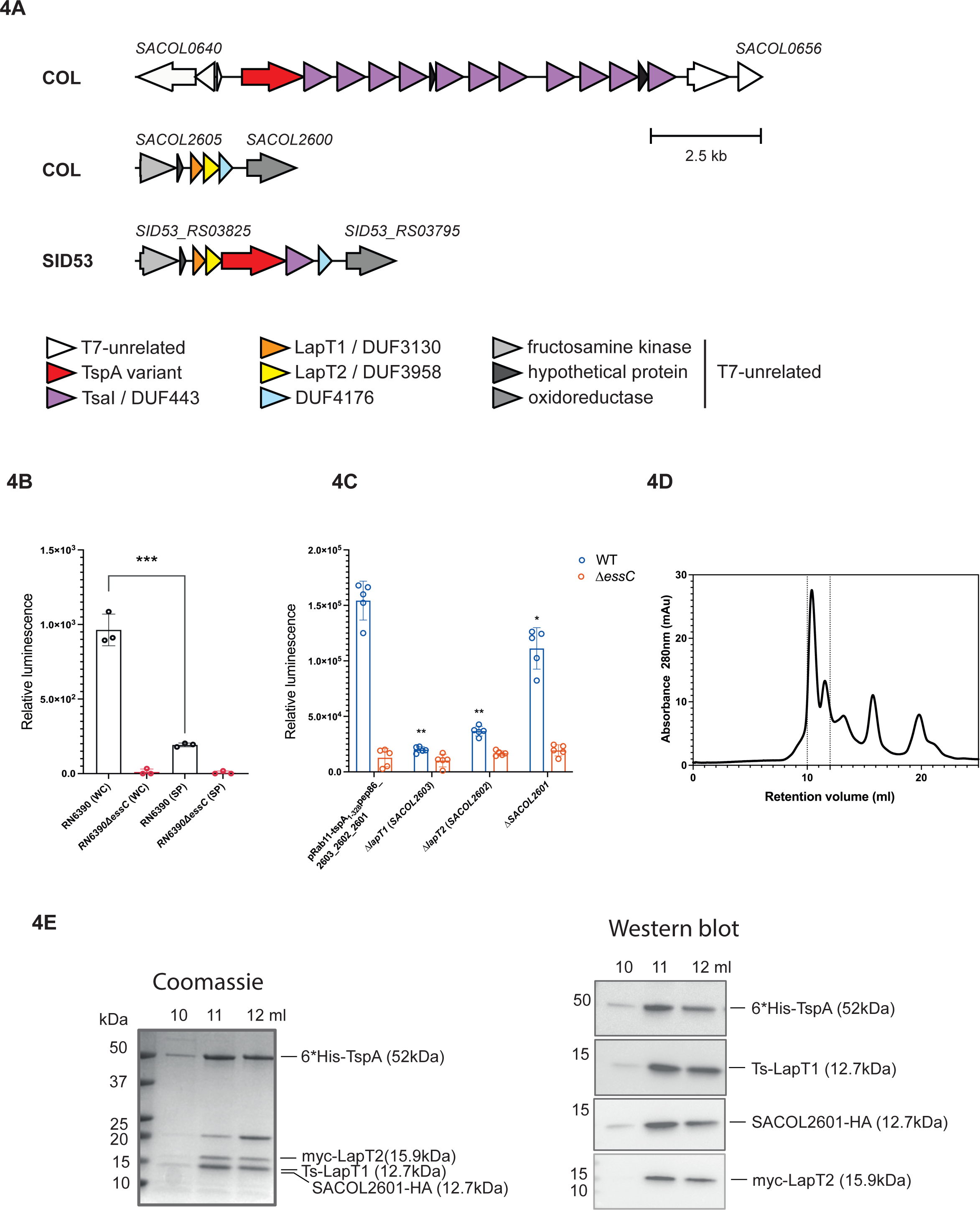
Identification of TspA secretion factors. A. Locus diagrams depicting the genetic neighbourhood of *tspA* (*SACOL0643*, red – top) and the T7SS-associated genes *SACOL2603*, *SACOL2602* and *SACOL2601* (orange, yellow and blue – middle) in strain COL. Bottom – the *SACOL2603* equivalent locus in strain SID53 has a *tspA2-tsaI2* insertion between the homologues of *SACOL2602* and *SACOL2601*. *SACOL2603* encodes a DUF3130/TIGR04917 family protein. B. COL wild-type or Δ*essC* strains carrying plasmid pRab11-tspA1-328-pep86_2603_2602_2601 were grown in 5 ml cultures at 37°C. At OD_600_ = 0.5 ATC was added to induce gene expression from pRab11. When cells reached OD_600_ = 2, samples corresponding to the whole cell culture (‘WC’) and the clarified culture medium (‘SP’) were prepared and 11S and furimazine were added to each. The data presented correspond to the mean luminescence readings of three biological replicates and error bars represent the SEM. Statistical significance for whole cell culture of the wild-type vs clarified culture medium was assessed with the Mann-Whitney test, with *** indicating p< 0.001. There was no significant difference between values for the whole cell vs clarified culture medium for the Δ*essC* strain. C. Wild-type and Δ*essC* variants of strain RN6390 were transformed with the indicated plasmids and cultured in 384-well plates at 34°C. Expression from pRab11 was induced by addition of ATC at 120 minutes, and luminescence was initiated by addition of furimazine and 11S at 260 minutes. Luminescence readings were taken at the 280 minute timepoint and were divided by OD_600_ readings to give the relative luminescence. Data represent the mean of five biological replicates each with 16 technical replicates. Error bars correspond to the SEM (n=5). The relative luminescence of RN6390 harbouring each plasmid containing the indicated individual gene deletion was compared with the relative luminescence signal of RN6390 harbouring pRab11-tspA_1-328_-pep86_2603-2602-2601 using the Mann-Whitney test, * p< 0.05; ** p<0.01. D. *E. coli* M15 cells carrying pQE70-_Tstrep_SACOL2603_myc_-2602-2601_HA_-_his_TspA were grown in LB medium. Following induction of gene expression with IPTG, complexes were purified by streptactin-affinity purification, and the resulting sample was analysed by size-exclusion chromatography (SEC). E. Fractions from within the dotted lines in (D) were subject to SDS-PAGE and analysed by coomassie staining (left panel) or by western blotting with antibodies against the tags on each component (right panel).

Genomic analysis previously identified a conserved module of three small genes, potentially implicated in T7 secretion, at a locus distant from both the *tspA* and *ess* gene clusters [4] (Fig. 4A, shaded orange, yellow and blue). In a handful of *S. aureus* strains, for example SID53, a second homologue of *tspA* is present at this locus, along with a DUF443-encoding immunity gene (Fig. 4A). However, in most well-studied strains, including COL, USA300 and RN6390 there is no *tspA* homologue or immunity gene at this locus, although the small genes are always present. This raised the possibility that the conserved cluster may be required for the biogenesis of TspA. Analysis of the encoded proteins indicates that SACOL2603 and SACOL2602 are predicted to be helical hairpin proteins belonging to the Lap1 (DUF3130) and Lap2 (DUF3958) families of WXG100-like export factors, respectively. Members of these protein families have been characterised in *Streptococcus intermedius* and shown to interact with the LXG domain of the lipid II phosphatase toxin, TelC [11, 26]. Following the nomenclature developed in *S. intermedius* and based on results described below we have renamed SACOL2603 and SACOL2602 LapT1 and LapT2, respectively (for <u>L</u>XG-associated <u>a</u>lpha helical <u>p</u>rotein for <u>T</u>spA). The third protein encoded at this locus, SACOL2601, has a DUF4176 domain. Genes encoding DUF4176 domains are frequently associated with T7SS loci [3] and the involvement of a DUF4176 domain-protein in LXG toxin secretion was recently determined in *S. intermedius* [27].

We therefore adapted the NanoBit assay to test the prediction that LapT1, LapT2 and SACOL2601 function in TspA secretion. We constructed a plasmid carrying *tspA_1-328_-pep86*, *SACOL2603*/*lapT1, SACOL2602*/*lapT2* and *SACOL2601*, avoiding issues arising from TspA toxicity by using a truncated version lacking the toxin domain but supplied with a C-terminal pep86 tag. Due to the slow growth kinetics of strain COL in 384-well plates we carried out assays in RN6390, whose sequence is identical to that of COL at the *tspA* and *SACOL2603* loci, except for a single (synonymous) nucleotide change in *tspA*. Following our observation of much lower EsxA-pep86 leakage into the supernatant at lower temperatures, we opted to conduct the secretion assays at 34°C.

After introduction of the plasmid into RN6390 wild-type and Δ*essC* strains, we measured luminescence in whole cell cultures, and in clarified culture supernatant following pelleting of cells (Fig. 4B). The presence of pep86-tagged TspA_NT_ in whole cell cultures was clearly detected, dependent on an active T7SS. However little or no TspA_NT_ was detected in the fractionated culture supernatant, confirming the previous finding that TspA is cell surface-associated rather than fully secreted [22]. These results also indicate that the TspA toxin domain (residues 329-469) is dispensable for cell surface attachment.

To determine whether any of LapT1, LapT2 and SACOL2601 are required to support secretion of TspA, we individually deleted each of the encoding genes from the plasmid and used the high throughput assay in whole cell cultures to assess export of pep86-tagged TspA_NT_. Fig. 4C indicates that in the absence of each gene, significantly less TspA_NT_-pep86 could be detected in whole cell cultures, while the cytoplasmic levels (deduced from measuring total luminescence per well following lysis) were not much affected (Fig S3B). We conclude that each of LapT1, LapT2 and SACOL2601 support TspA secretion. It should be noted the chromosomal copy of the *SACOL2601-2603* gene cluster is present in these strains and may account for some of the residual secretion seen from the plasmid deletion constructs.

Previous work has reported that LXG proteins form a stable pre-secretion complex with their cognate helical hairpin secretion partners [23, 24, 26, 27]. To investigate whether any of LapT1, LapT2 or SACOL2601 interact with TspA, we co-produced all four proteins in *E. coli*. The recombinant proteins carried an N-terminal twinstrep tag on LapT1 for purification, and a C-terminal HA, N-terminal myc and N-terminal his tag on SACOL2601, LapT2 and TspA, respectively, for detection. Complexes were purified by streptactin affinity chromatography and analysed by size exclusion chromatography (SEC; Fig. 4D) and both SDS PAGE and western blotting (Fig. 4E). All four proteins co-eluted in the SEC main peak confirming that they exist as a tetrameric complex. Taken together our results indicate that the *SACOL2601-2603* gene cluster is required for the biogenesis of TspA.

## Discussion

Here we report a novel, sensitive assay for monitoring protein secretion by the T7SS in *S. aureus*, using the NanoBit split luciferase system. The 11 amino acid pep86 fragment of NanoLuc is small enough to be compatible with secretion when fused to the WXG100 protein EsxA, or to a larger LXG protein substrate. As the presence of luminescence from secreted proteins can be measured directly in unprocessed cell cultures, this enables high throughput screening of large numbers of strains and growth conditions.

Most studies of the *S. aureus* T7SS to date have focussed on secretion by *essC-1* strains, using a handful of isolates. Strains exhibit high genetic diversity at their T7 loci [14], but experimental investigation of *essC-2* - *essC-4* variant strains has previously been constrained by low activity in lab conditions, and the lack of a sensitive secretion assay [33]. Using the NanoBit assay we found substantial variability in the secretion efficiencies of such strains, and their temperature dependence. In particular, the EMRSA15 T7SS (an *essC-4* variant strain), which was inactive at 37°C, exhibited a significant level of secretion at 30°C and 34°C suggesting a role for temperature in T7SSb regulation. Interestingly, *ess* transcription and T7 secretion were previously shown to be activated by reducing membrane fluidity, which was achieved by treating cells with *cis*-unsaturated fatty acids [38, 39]. Temperature-induced changes in membrane fluidity may potentially account for the differences in secretion observed here.

A number of pathways have been implicated in transcriptional regulation of the *S. aureus* T7SS, including the SaeRS and ArlRS two-component systems, the Agr quorum-sensing system and the alternative sigma factor SigB [19, 38, 40–43]. However, the complex regulatory picture is far from complete and key questions around how, why and when the secretion system is expressed remain unanswered. Moreover, additional factors may activate secretion at the post transcriptional level, for example the presence of haemin has been linked to post-transcriptional activation of the T7SS in some strains [33, 44]. The availability of a high throughput assay will facilitate screening of a range of conditions, paving the way for a much more detailed analysis of T7 regulation than has been possible to date.

We employed the NanoBit assay to explore the secretion of the surface-attached LXG toxin, TspA. While other LXG toxins characterised to date are encoded alongside accessory proteins essential for their biogenesis [24, 26, 27], secretion partners for TspA had not been identified. A conserved cluster of three genes remote from the *tspA* locus had previously been bioinformatically linked to the T7SS [4]. We showed that each of these genes supports secretion of a TspA-pep86-fusion suggesting that their genomic conservation is because they are required for TspA biogenesis. This conclusion was further supported by co-purification experiments, where tagged variants of the three proteins formed a complex with TspA. Two of these genes, *SACOL2603* and *SACOL2602* code for helical hairpin proteins of the DUF3130 and DUF3958 families, respectively. The LXG toxins, TelC and TelD, from *S. intermedius* both require proteins from each of these families for secretion, with Lap1, being a DUF3130 protein and Lap2 a DUF3958. In each case the Lap1 and Lap2 proteins were shown to interact with the LXG domain of their cognate toxin partners. In keeping with this nomenclature, we have renamed SACOL2603 as LapT1 and SACOL2602 as LapT2. Klein *et al*. [26] identified a conserved FxxxD motif close to the C-terminus of LapD1, showing that individual alanine substitutions of either conserved residue was sufficient to prevent TelD secretion. LapD1 and LapT1 share only limited sequence similarity (Fig. S4) however an FxxxD motif is present at the same position in both proteins. This motif is conserved across LapT1 homologues in other staphylococci and in enterococci. It remains to be determined whether this serves as a secretion motif for TspA export.

The third protein gene, *SACOL2601*, codes for a DUF4176 protein. Interestingly, proteins of this family are also encoded in the *S. intermedius telA*, *telB* and *telD* gene clusters, but not *telC*. While the homologue encoded at the *telA* locus was shown to be essential for TelA secretion, it was not required for TelC export [27]. It is not clear whether TelC requires a DUF4176 protein encoded by a different cluster, or whether its secretion occurs independently of such a protein. In addition to TspA, a variant-specific LXG protein coding gene is found at the *ess* loci of *essC-1*, essC*-2*, *essC-3* and *essC-4 S. aureus* strains [4]. However, no DUF4176-encoding genes are found at any *ess* loci, suggesting either they are not universally required, or that a single DUF4176 protein might function in secretion of multiple substrates. Further work would be required to investigate this.

In conclusion we describe a novel, sensitive assay to assess secretion of T7 substrate proteins in *S. aureus*. We anticipate this approach should be broadly applicable for the study of type VII secretion across a range of organisms.

## Acknowledgements

This study was supported by Wellcome Trust Investigator awards 10183/Z/15/Z and 224151/Z/21/Z to TP, a China Scholarship Council PhD studentship to YY and a Newcastle University research scholarship to AAS. We thank Ben Berks and Xiaolong Liu for advice and reagents for the NanoBit assay, Ian Collinson for plasmid pBAD-_6H_11S for purification of 11S, José Penadés and Ross Fitzgerald for *S. aureus* strain JP5347, and Sharon Peacock for strain EMRSA15.

## Data, code and materials

The datasets supporting this article have been uploaded to FigShare (https://doi.org/10.6084/m9.figshare.25368598).

## Notes

### Competing Interest Statement

The authors have declared no competing interest.

### Summary of Updates

Minor corrections - Figure S4 has been corrected

