## Supplementary information for "A high throughput assay provides novel insight into type VII secretion in *Staphylococcus aureus*"

Table S1. **Oligonucleotides used in this study**

| Name | Sequence | Template | Construct |
|---|---|---|---|
| pRab11_fwd | GAATTCACTGGCCGTCGTTTTAC | pRab11 | pRab11-esxApe86 |
| pRab11_rev | GGTACCATCATACTCTATCAATGATAG | pRab11 | pRab11-esxApep86 |
| esxA_fwd | gatagagtatgatggtaccaggaggtttctagttATGGCAATGATTAAGATGAG | COL gDNA | pRab11-esxApep86 |
| esxA_rev | aaacgacggccagtgaattcttagctaatttttttaaacaggcgccagccgctcacgccggagccTTGCAAACCGAAATTATTAG | COL gDNA | pRab11-esxApep86 |
| TrxA_fwd | tgatagagtatgatggtaccaggaggtttctagttatggtgagcggctggcgcctgtttaaaaaaattagcggctccggcGCAATCGTAAAAGTAACAG | COL gDNA | pRab11-pep86trxA |
| TrxA_rev | aaacgacggccagtgaattcTTATAAATGTTTATCTAAAACTTCAGC | COL gDNA | pRab11-pep86trxA |
| EssCdA1 A | AGTACTGAATTCGTATGATG | | All pIMAYessC constructs |
| EssC398dA2 B | CTATTGAATTAACAATTTATGCATTGTCTTTGCCTCAG | MRSA252/10.1252.X gDNA | pIMAYessC-ST398/pIMAYessC-MRSA252 |
| EssC398dB2 C | ATGCATAAATTGTTAATTCAATAGGGAGGACAATTATG | 10.1252.X gDNA | pIMAYessC-ST398 |
| EssC398dB1 D | ATTGACGAATTCAAATTGGTGCG | 10.1252.X gDNA | pIMAYessC-ST398 |
| EssC252dB2 | ATGCATAAATTGTTAATTCAATAGGGAGGACAATTATG | MRSA252 gDNA | pIMAYessC-MRSA252 |
| EssC252dB1 | ATTGACGAATTCCTTCTTCGCATC | MRSA252 gDNA | pIMAYessC-MRSA252 |
| EssCEMRSA15dA2 | TTATTCAAATAACAATTTATGCATTGTCTTTGCCTCAG | EMRSA15 gDNA | pIMAYessC-EMRSA15 |
| EssCEMRSA15dB1 | CTTATAGAATTCAAACAATTTTCGTAATTTAGC | EMRSA15 gDNA | pIMAYessC-EMRSA15 |
| EssCEMRSA15dB2 | ATGCATAAATTGTTATTTGAATAAAAGGAGAGTATTATG | EMRSA15 gDNA | pIMAYessC-EMRSA15 |
| pIMAY-Z_fwd | CGGCCGCCACCGCGGTGGAG | pIMAY-Z | pIMAY-Z_esxApep86 |
| pIMAY-Z_rev | CGATATCAAGCTTATCGATACCGTCGACCTCGAGGGGG | pIMAY-Z | pIMAY-Z_esxApep86 |
| esxA stop codon up-700_fwd | tatcgataagcttgatatcgAGTATACGCGCCGGTGTCTTTATTC | COL gDNA | pIMAY-Z_esxApep86 |
| esxA stop codon up-700_rev | TTGCAAACCGAAATTATTAGAAAGTTGTTG | COL gDNA | pIMAY-Z_esxApep86 |
| esxA stop codon down-700_fwd | ctaataatttcggtttgcaaggctccggcgtgagcggctggcgcctgtttaaaaaaattagcTAAGCATTCTGAAATTGGCAAAG | COL gDNA | pIMAY-Z_esxApep86 |
| esxA stop codon down-700_rev | ctccaccgcggtggcggccgGGGAAGTCGTTAATTGGATTTAATAAG | COL gDNA | pIMAY-Z_esxApep86 |

| | | | |
|---|---|---|---|
| YPAS5_pRab11_fwd | GGTACCGTTAACAGATCTG | pRab11 | pRab11-tspA$_{1-328}$-pep86-SACOL2603_SACOL2602_SACOL2601 |
| YPAS2_pRab11_R | CTTTTTCAAAATCATACTCTATCAATGATAGAG | pRab11 | pRab11-tspA$_{1-328}$-pep86-SACOL2603_SACOL2602_SACOL2601 |
| YPAS3 rbs-TspA1-328_fwd | AGAGTATGATTTTGAAAAAGGAGCATGC | COL gDNA | pRab11-tspA$_{1-328}$-pep86-SACOL2603_SACOL2602_SACOL2601 |
| YPAS4 rbs-TspA1-328_rev | TCAGCTAATTTTTTTAAACAGGCGCCAGCCGCTCACTCCGCCTTTTGCAAATTTAC | COL gDNA | pRab11-tspA$_{1-328}$-pep86-SACOL2603_SACOL2602_SACOL2601 |
| YPAS6 rbs-SACOL2603-1_fwd | GTGAGCGGCTGGCGCCTGTTTAAAAAAATTAGCTGATTAAATGGAGAGAGGTGTAAATG | COL gDNA | pRab11-tspA$_{1-328}$-pep86-SACOL2603_SACOL2602_SACOL2601 |
| YPAS7 rbs-SACOL2603-1_rev | TCAGATCTGTTAACGGTACCTTATTGTTGTGTAACTTCTCC | COL gDNA | pRab11-tspA$_{1-328}$-pep86-SACOL2603_SACOL2602_SACOL2601 |
| AS2-2601_F | CTCAGTACAATCTGCTCTG | pRab11-tspA$_{1-328}$-pep86-SACOL2603-1 | pRab11-tspA$_{1-328}$-pep86-SACOL2603-SACOL2602 |
| AS2-2601_R | TTTTGCGAACCTTCTTTTAAATAAATAATG | pRab11-tspA$_{1-328}$-pep86-SACOL2603-1 | pRab11-tspA$_{1-328}$-pep86-SACOL2603-SACOL2602 |
| AS2-2602_F | AAGAAGAGATGATTGACTTG | pRab11-tspA$_{1-328}$-pep86-SACOL2603-1 | pRab11-tspA$_{1-328}$-pep86-SACOL2603-SACOL2601 |
| AS2-2602_R | CGACATCATTTCACCACG | pRab11-tspA$_{1-328}$-pep86-SACOL2603-1 | pRab11-tspA$_{1-328}$-pep86-SACOL2603-SACOL2601 |
| As2-2603-F | AATGATTGGTAATGAAATTGG | pRab11-tspA$_{1-328}$-pep86-SACOL2603-1 | pRab11-tspA$_{1-328}$-pep86-SACOL2602-SACOL2601 |

| | | | |
|---|---|---|---|
| As2-2603-R | TCATTTACACCTCTCTCC | pRab11-tspA$_{1-328}$-pep86-SACOL2603-1 | pRab11-tspA$_{1-328}$-pep86-SACOL2602-SACOL2601 |
| pQE70_fwd | TAAGCTTAATTAGCTGAGC | pQE70 | pQE70-$_{Tstrep}$SACOL2603-$_{myc}$2602-2601$_{HA\text{-}his}$TspA |
| pQE70_rev | TGAATTCTGTGTGAAATTGTTATC | pQE70 | pQE70-$_{Tstrep}$SACOL2603-$_{myc}$2602-2601$_{HA\text{-}his}$TspA |
| tspA_fwd | acaatttcacacagaattcaaaagaggagaaaatgcatcaccatcaccatcacAGTATTGACATGTATTTAGACAGATCTC | COL gDNA | pQE70-$_{Tstrep}$SACOL2603-$_{myc}$2602-2601$_{HA\text{-}his}$TspA |
| tspA_rev | TCACCACGCCCATTTCATTG | COL gDNA | pQE70-$_{Tstrep}$SACOL2603-$_{myc}$2602-2601$_{HA\text{-}his}$TspA |
| Ts_fwd | caatgaaatgggcgtggtgaaaagaggagaaattaagcatgAGTGCTTGGAGTCATCCAC | pQE70-Ts | pQE70-$_{Tstrep}$SACOL2603-$_{myc}$2602-2601$_{HA\text{-}his}$TspA |
| Ts_rev | ctgtactcatCTTCTCAAATTGTGGATGAGAC | pQE70-Ts | pQE70-$_{Tstrep}$SACOL2603-$_{myc}$2602-2601$_{HA\text{-}his}$TspA |
| 2603_fwd | atttgagaagATGAGTACAGTTCAAAGTG | COL gDNA | pQE70-$_{Tstrep}$SACOL2603-$_{myc}$2602-2601$_{HA\text{-}his}$TspA |
| 2603_rev | TTAATTTTTACCAATTTCATTACCAATC | COL gDNA | pQE70-$_{Tstrep}$SACOL2603-$_{myc}$2602-2601$_{HA\text{-}his}$TspA |
| myc2602-2601HA_fwd | aatgaaattggtaaaaattaaaaagaggagaaaatggaacaaaaactcatctcagaagaggatctgTCGAATAAACTGGATGAAATC | COL gDNA | pQE70-$_{Tstrep}$SACOL2603-$_{myc}$2602-2601$_{HA\text{-}his}$TspA |
| myc2602-2601HA_rev | agctcagctaattaagcttaagcgtaatctggaacatcgtatgggtaTTGTTGTGTAACTTCTCC | COL gDNA | pQE70-$_{Tstrep}$SACOL2603-$_{myc}$2602-2601$_{HA\text{-}his}$TspA |

Figure S1

A. Growth curves for cultures of the indicated wild-type and Δ*essC* strains, each carrying pRab11-esxApep86. 50 µl cultures (three biological replicates, each with 16 technical replicates) were grown in 384-well plates at 37°C. ATC was added at 180 min, furimazine and 11S were added at 220 min (silver dotted lines) for all *S. aureus* strains except COL and its *essC* derivative. Here, ATC was added at 320 min, furimazine and 11S were added at 380 min (pink dotted lines). In each case $OD_{600}$ readings were taken every ten minutes and mean $OD_{600}$ values are plotted.

B. Following the initiation of luminescence in the cultures in (A), luminescence readings were taken at ten minute intervals, and mean values are plotted.

C. Cultures were grown as in (A) and at the time point corresponding to the peak value in Fig. S1B, were diluted either 1:2 or 1:4 in TBS with 50 µg/ml lysostaphin. Furimazine and 11S were added and luminescence readings taken after 20 minutes. Luminescence values were divided by the $OD_{600}$ at that timepoint to give the relative luminescence. Error bars correspond to the SEM (n=3). WT – wild-type.

Figure S2

A. Growth curves for cultures of the indicated wild-type and Δ*essC* strains, each carrying pRab11-esxApep86. 50 µl cultures (three biological replicates, each with 16 technical replicates) were grown in 384-well plates at 30°C (left) or 34°C (right). ATC was added 220 minutes (34°C cultures), or 240 minutes (30°C cultures), furimazine and 11S were added at 280 minutes (34°C cultures) or 300 minutes (30°C cultures), and $OD_{600}$ readings were taken every ten minutes. Mean $OD_{600}$ values are plotted.

B. Following the initiation of luminescence in the cultures in (A), luminescence readings were taken at ten minute intervals, and mean values are plotted.

C. Cultures were grown as in (A) and at the time point corresponding to the peak value in Fig. S2B, were diluted either 1:2 or 1:4 in TBS with 50 µg/ml lysostaphin. Furimazine and 11S were added and luminescence readings taken after 20 minutes. Luminescence values were divided by the $OD_{600}$ at that timepoint to give the relative luminescence. Error bars correspond to the SEM (n=3). WT – wild-type.

Figure S3.

A. Growth curves for cultures of RN6390 and its Δ*essC* derivative carrying the indicated plasmids. 50 µl cultures (five biological replicates, each with 16 technical replicates) were grown in 384-well plates at 34°C. $OD_{600}$ readings were taken every ten minutes and mean values are plotted.

B. Cultures were grown as in (A). Expression of encoded genes from pRAB11 were induced by addition of ATC at 120 minutes, and luminescence was initiated by addition of furimazine and 11S at 260 minutes. Luminescence readings were taken at the 280 minute timepoint and-were divided by $OD_{600}$

readings to give the relative luminescence. Data represent the mean of five biological replicates with 64 cultures per replicate. Error bars correspond to the SEM (n=5). WT – wild-type.

Figure S4.

Alignment of LapT1 (SACOL2603) from *S. aureus* with LapD1 from *S. intermedius*. The two proteins share 20.9% identity. The FxxxD motif is highlighted in yellow.

Figure S5.

Uncropped blots and gels for all experimental data.

S1A

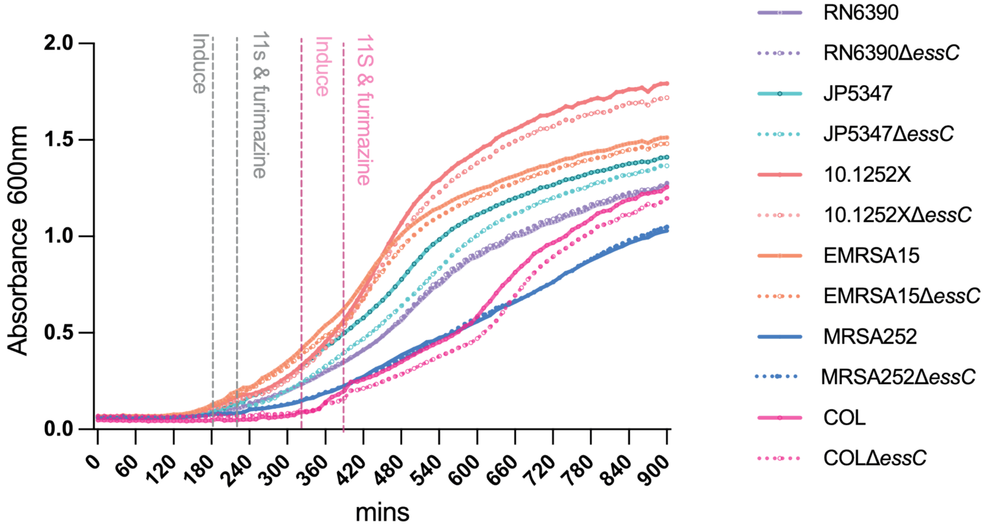
Induce
11s & furimazine
Induce
11S & furimazine
Absorbance 600nm
2.0
1.5
1.0
0.5
0.0
0 60 120 180 240 300 360 420 480 540 600 660 720 780 840 900
mins
RN6390
RN6390ΔessC
JP5347
JP5347ΔessC
10.1252X
10.1252XΔessC
EMRSA15
EMRSA15ΔessC
MRSA252
MRSA252ΔessC
COL
COLΔessC

S1B

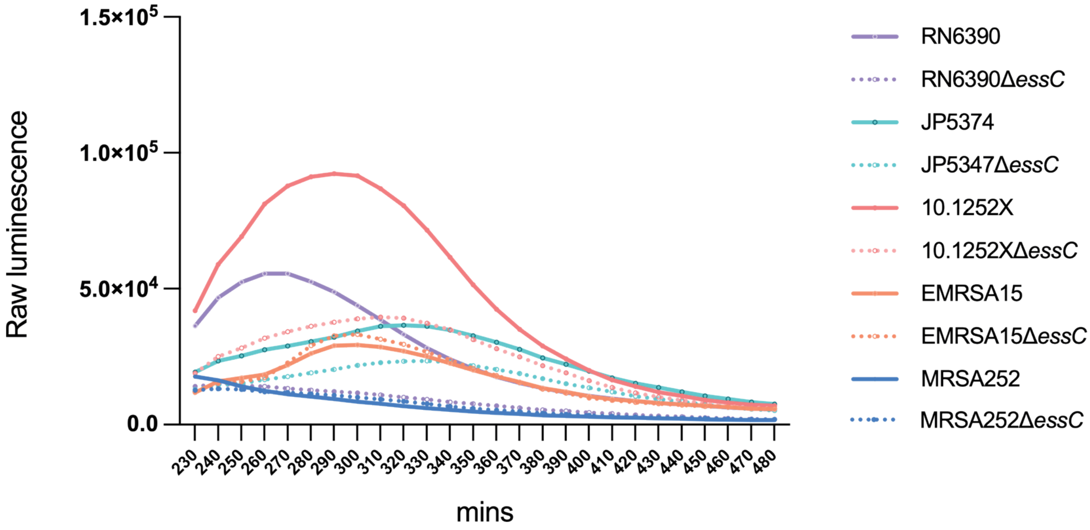
Raw luminescence
1.5×10^5
1.0×10^5
5.0×10^4
0.0
230 240 250 260 270 280 290 300 310 320 330 340 350 360 370 380 390 400 410 420 430 440 450 460 470 480
mins
RN6390
RN6390ΔessC
JP5374
JP5347ΔessC
10.1252X
10.1252XΔessC
EMRSA15
EMRSA15ΔessC
MRSA252
MRSA252ΔessC

S1C

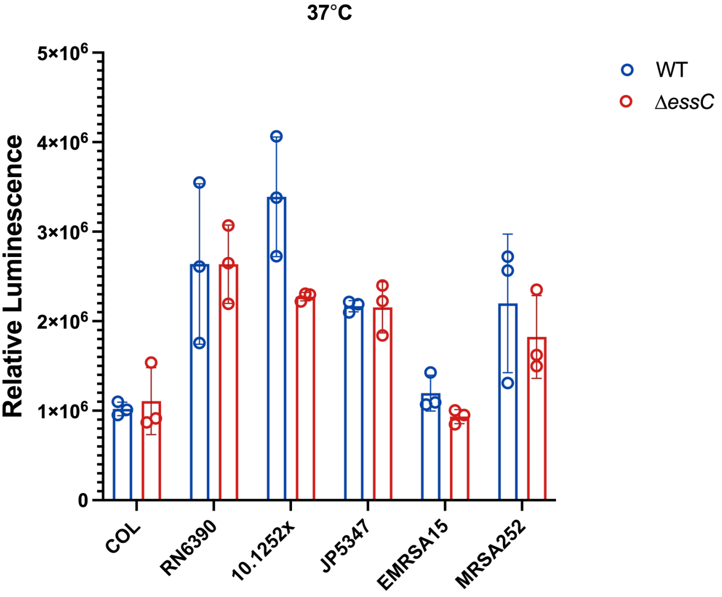
37°C
Relative Luminescence
5×10^6
4×10^6
3×10^6
2×10^6
1×10^6
0
COL
RN6390
10.1252x
JP5347
EMRSA15
MRSA252
WT
ΔessC

## S2A

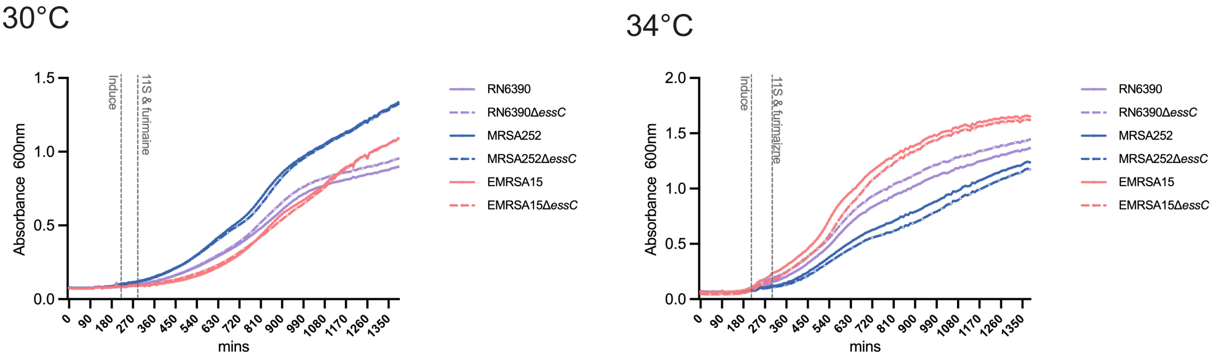
30°C
34°C
Induce
11S & furimaine
11S & furimaizne
Absorbance 600nm
mins
0.0
0.5
1.0
1.5
2.0
0
90
180
270
360
450
540
630
720
810
900
990
1080
1170
1260
1350
RN6390
RN6390ΔessC
MRSA252
MRSA252ΔessC
EMRSA15
EMRSA15ΔessC

## S2B

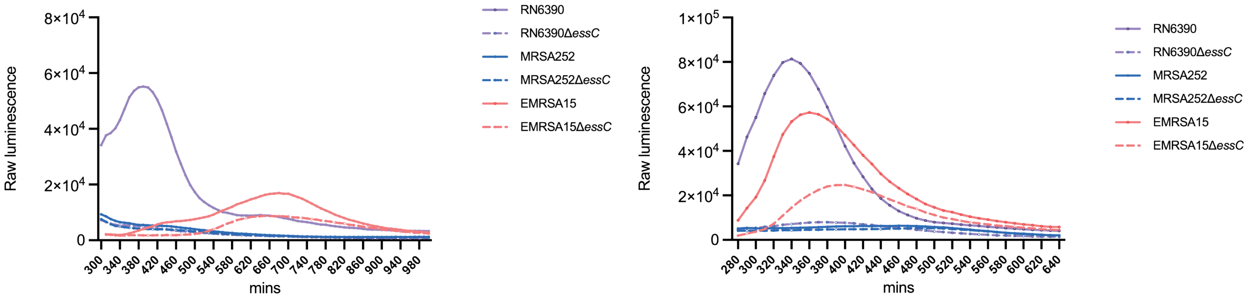
Raw luminescence
mins
0
2×10^4
4×10^4
6×10^4
8×10^4
1×10^5
300
340
380
420
460
500
540
580
620
660
700
740
780
820
860
900
940
980
280
320
360
400
440
480
520
560
600
640
RN6390
RN6390ΔessC
MRSA252
MRSA252ΔessC
EMRSA15
EMRSA15ΔessC

## S2C

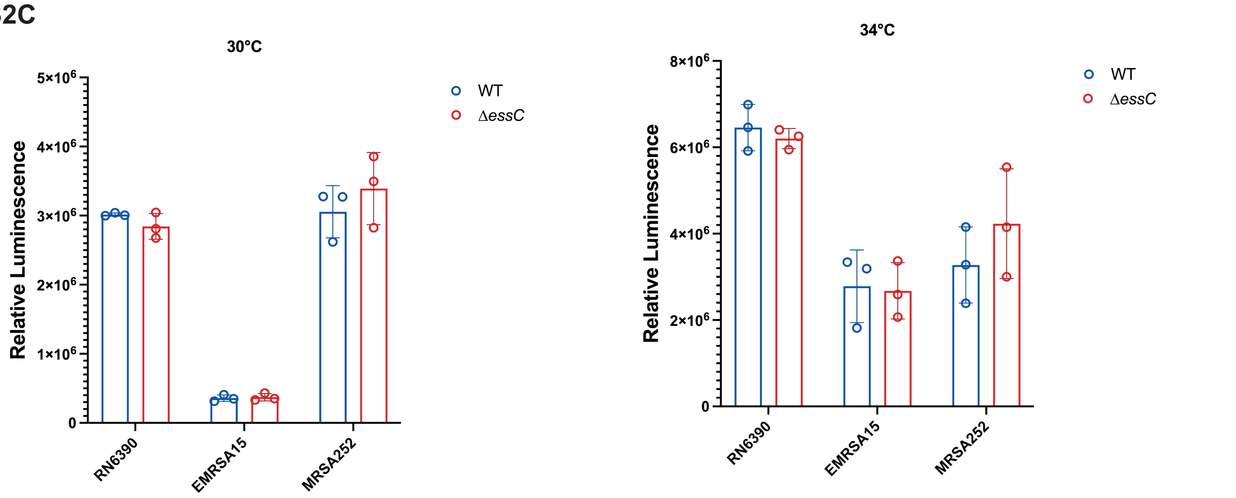
30°C
34°C
Relative Luminescence
0
1×10^6
2×10^6
3×10^6
4×10^6
5×10^6
6×10^6
8×10^6
RN6390
EMRSA15
MRSA252
WT
ΔessC

## S3A

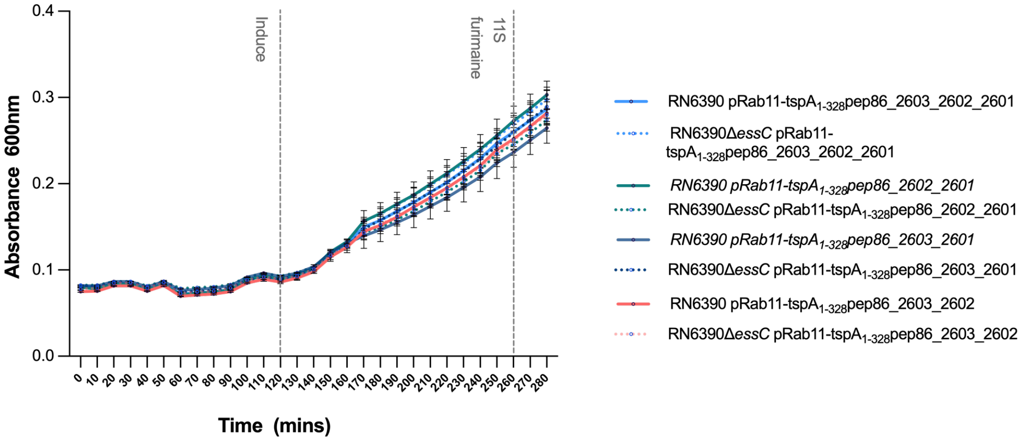
Absorbance 600nm
0.0
0.1
0.2
0.3
0.4
Time (mins)
0 10 20 30 40 50 60 70 80 90 100 110 120 130 140 150 160 170 180 190 200 210 220 230 240 250 260 270 280
Induce
11S furimaine
RN6390 pRab11-tspA1-328pep86_2603_2602_2601
RN6390ΔessC pRab11-tspA1-328pep86_2603_2602_2601
RN6390 pRab11-tspA1-328pep86_2602_2601
RN6390ΔessC pRab11-tspA1-328pep86_2602_2601
RN6390 pRab11-tspA1-328pep86_2603_2601
RN6390ΔessC pRab11-tspA1-328pep86_2603_2601
RN6390 pRab11-tspA1-328pep86_2603_2602
RN6390ΔessC pRab11-tspA1-328pep86_2603_2602

## S3B

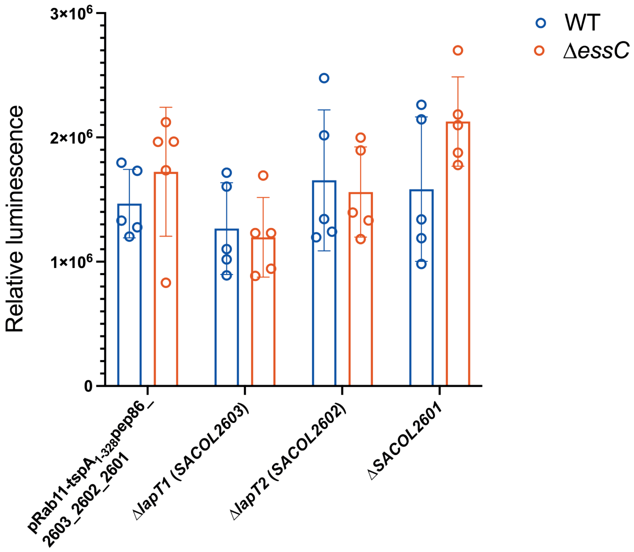
Relative luminescence
0
1×10⁶
2×10⁶
3×10⁶
WT
ΔessC
pRab11-tspA1-328pep86_2603_2602_2601
ΔlapT1 (SACOL2603)
ΔlapT2 (SACOL2602)
ΔSACOL2601

```
LapT1  MMSTVQSDIFKTNSASSSIKSAVETCNNVSKPDKDESTTVSGNNNAHSVIDDLMSKNQSVAEAIRTASDNIQKVGEAFDQTDVMIGNEIGKN------------------------    92
LapD1   MGNIKSEAGVAQSVASGIMTGAGSISDVGQAHPDEQSRYSGNDIAKDKIMSESNYGELLSNVLQRFIHLIHTTAAEFVGVDQHLAHEIEKSSSTAQATSRGVRNPNFVPDRSLFK    115
        *..::*:   ::*.:*.* :.. : .:*.: . **.:  ***: *:. * .  . .: :::.::   . *:...  *  .*  :.:** *.
```

Fig. S5

Fig.1C top panel

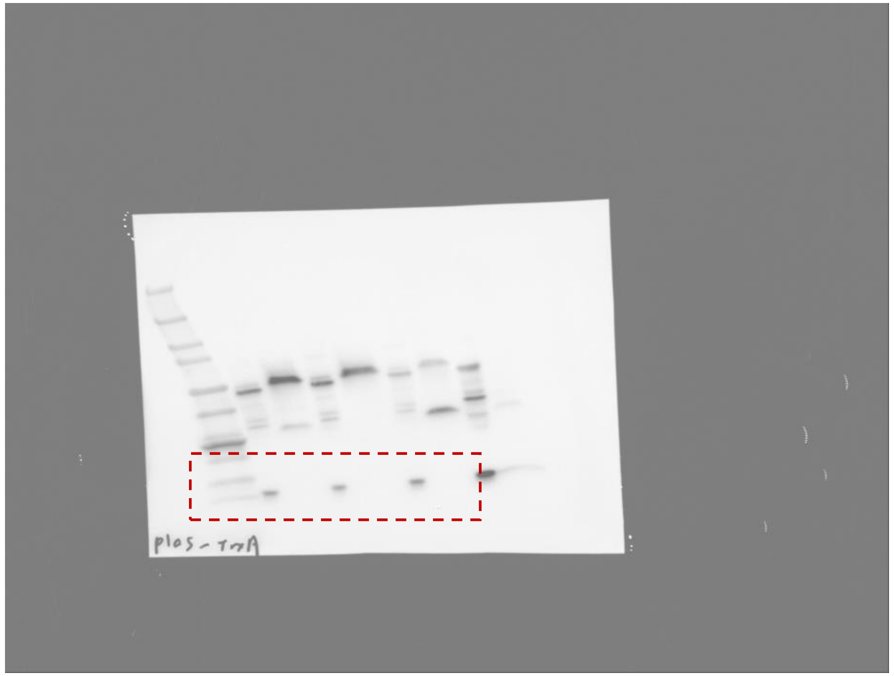


Fig.1C lower panel

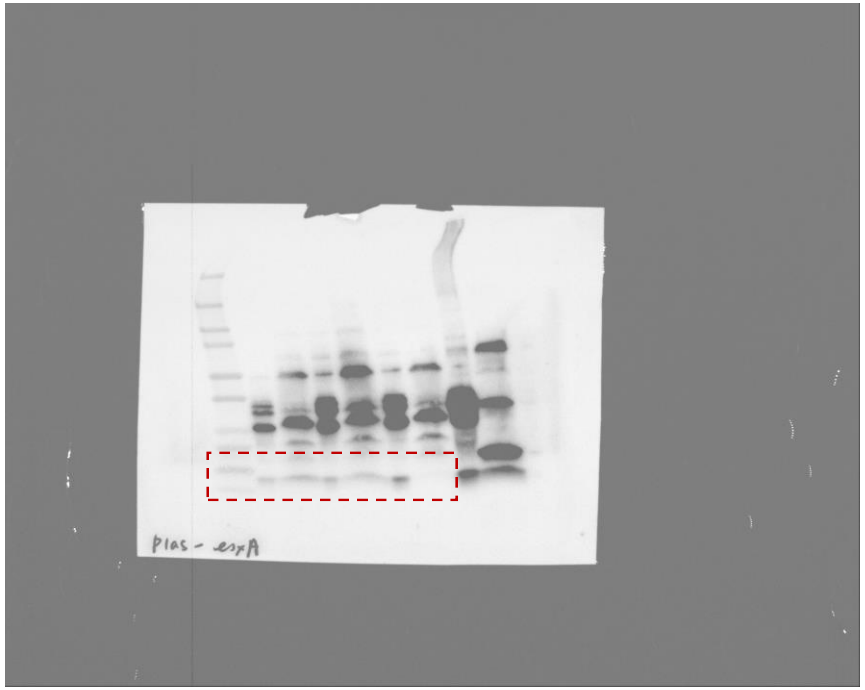

Fig.1E top panel

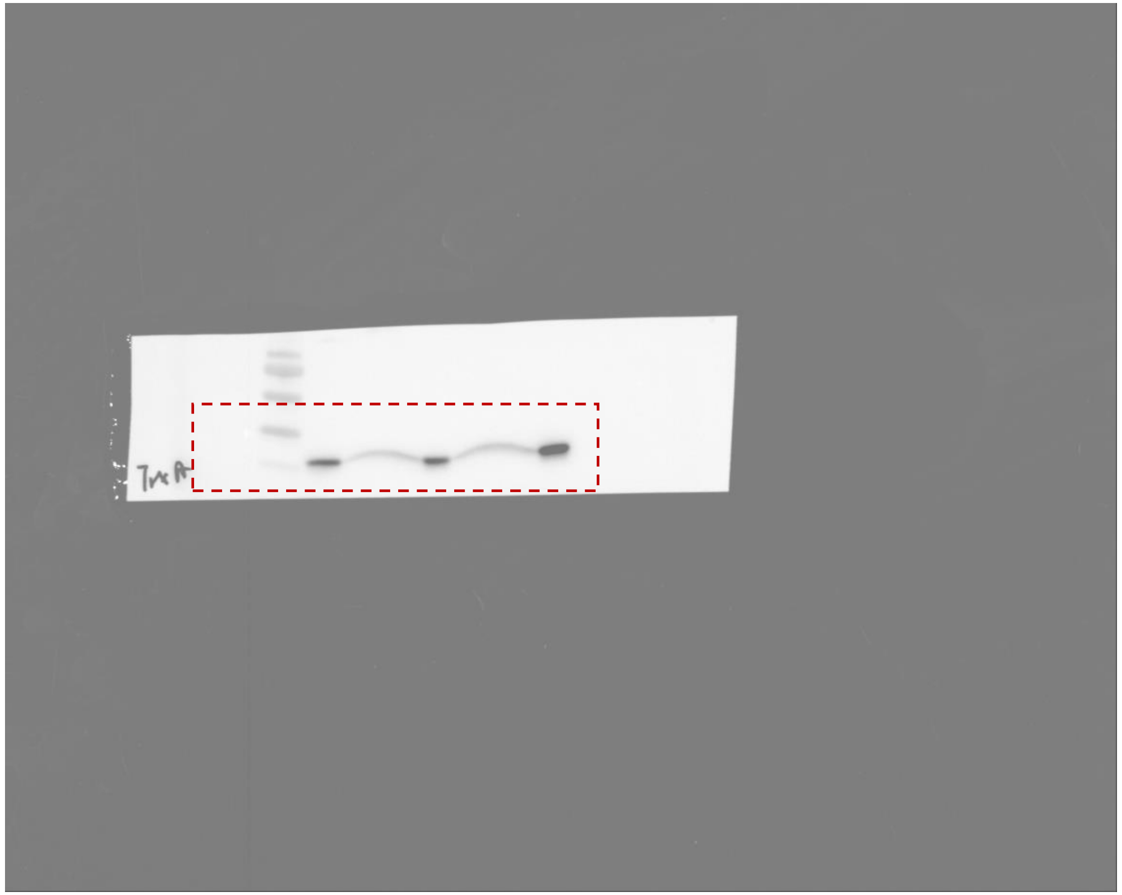

Fig.1E lower panel

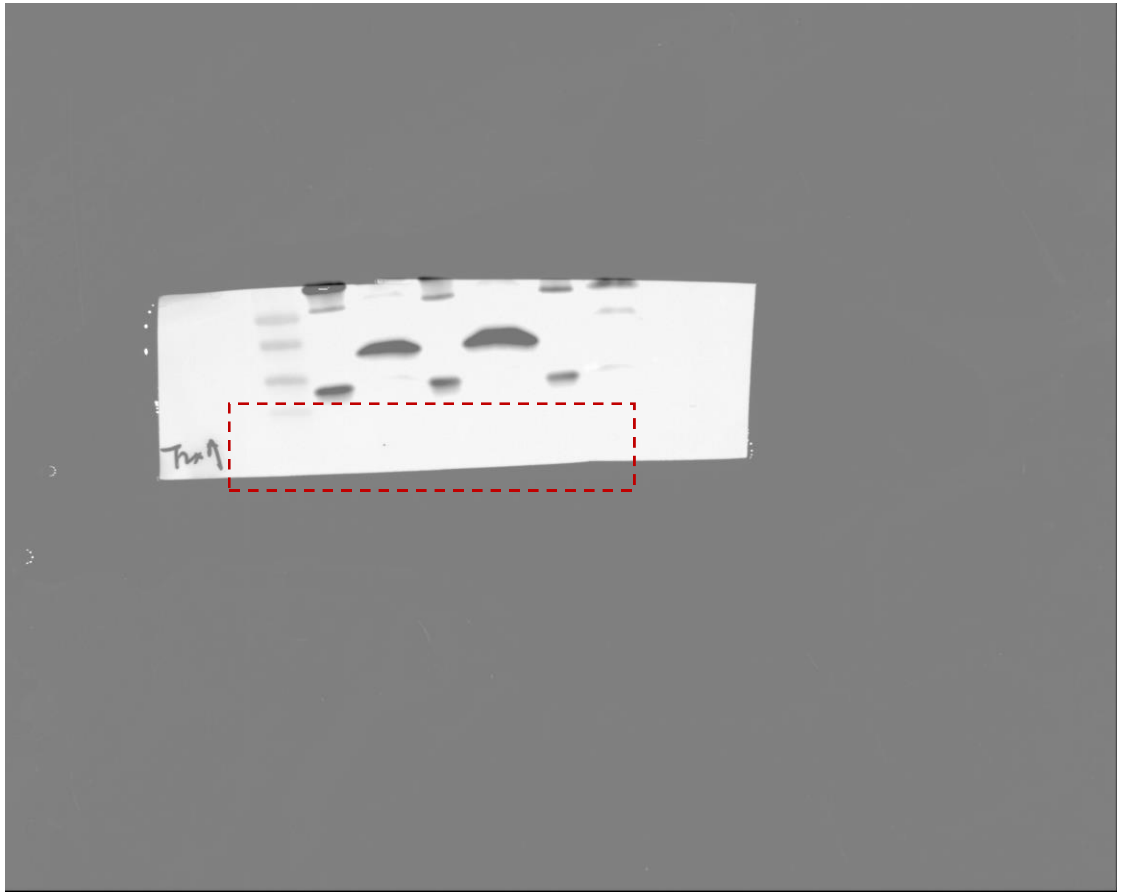

Fig. 2A top panel

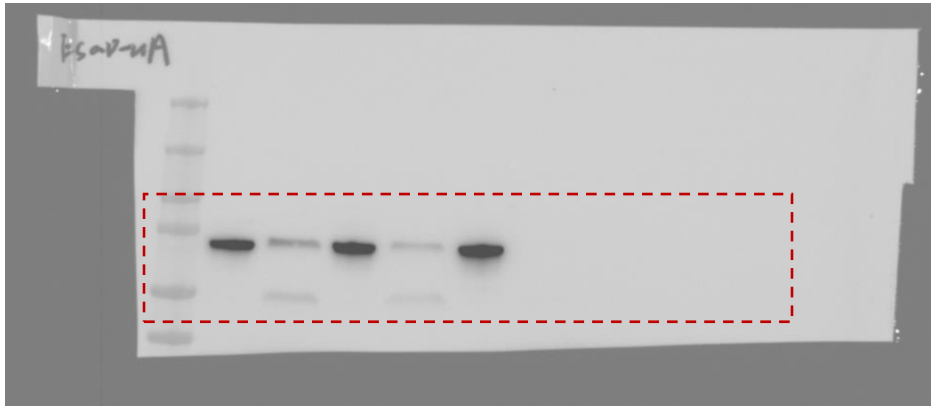

Fig. 2A middle panel

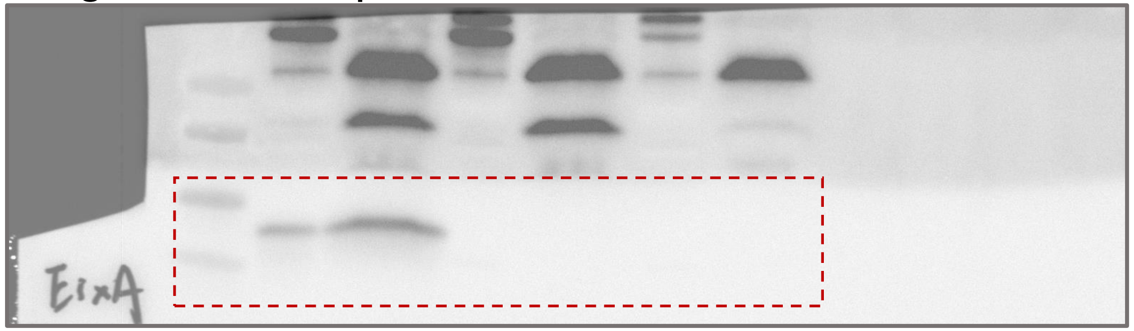

Fig. 2A lower panel

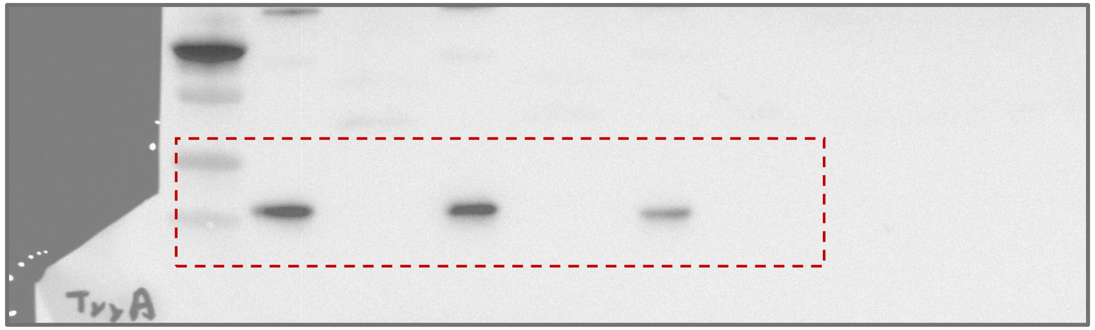

Fig. 4E Coomassie

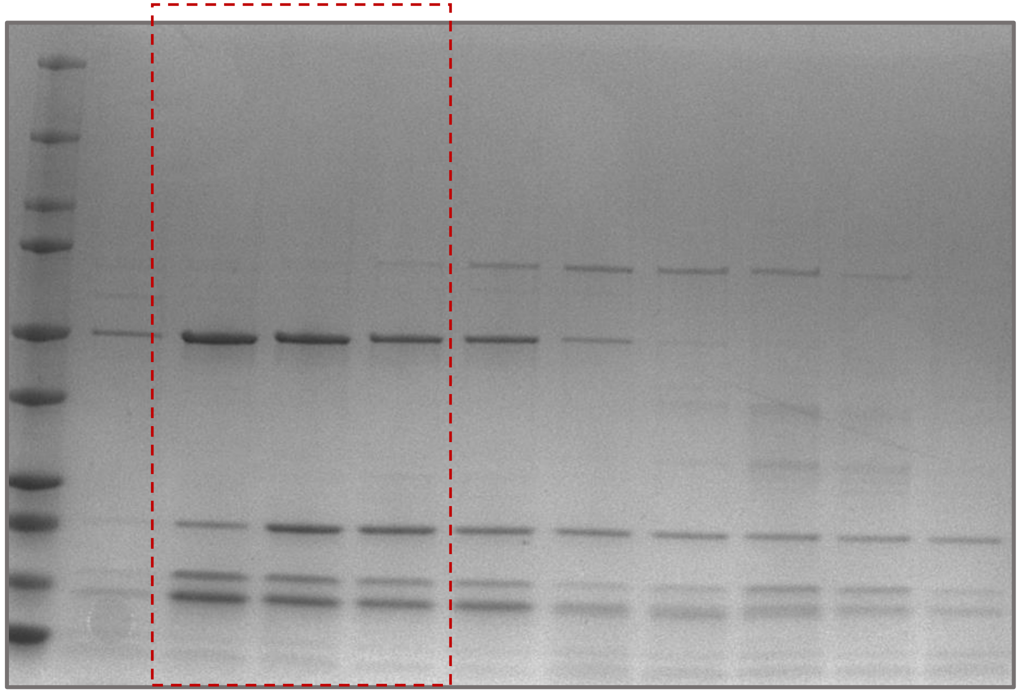

Fig. 4E blot-1

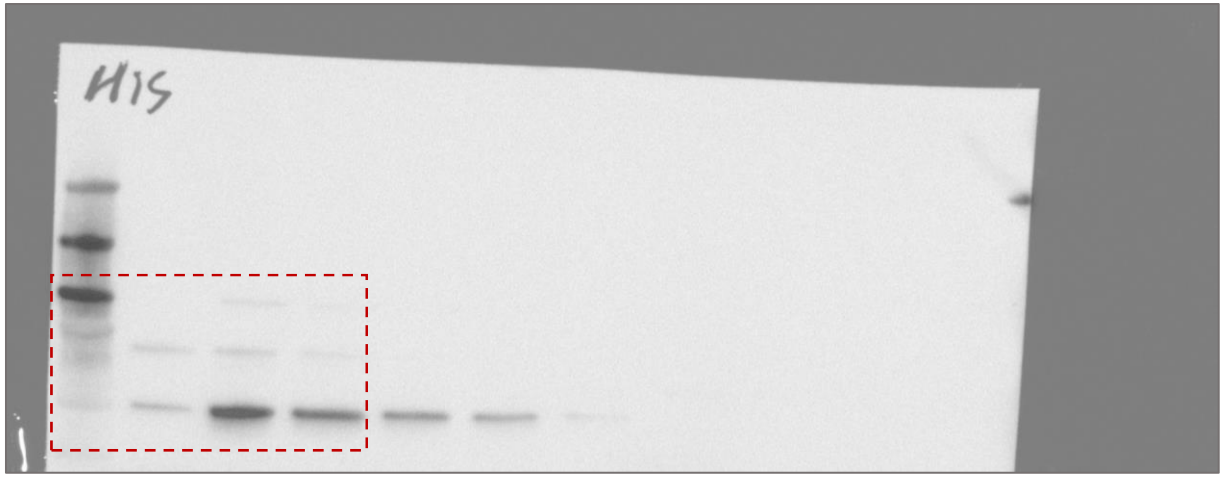


Fig. 4E blot-2

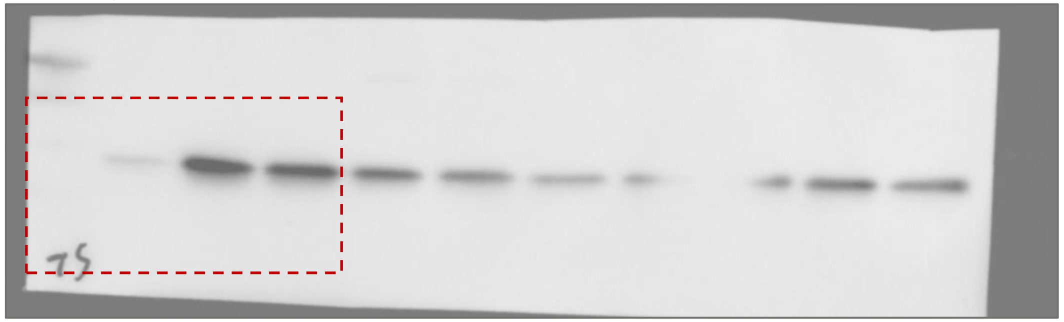

Fig. 4E blot-3

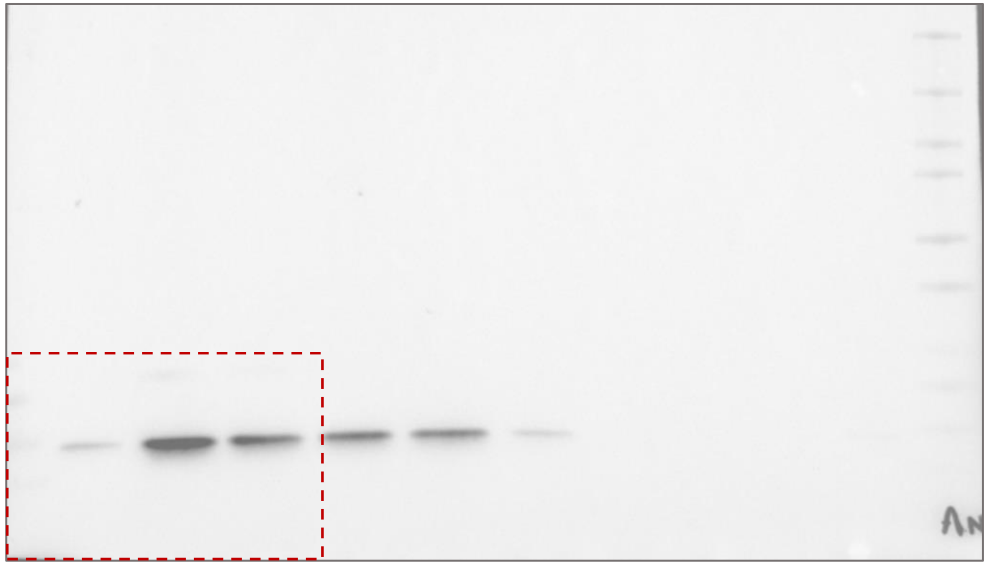

Fig. 4E blot-4

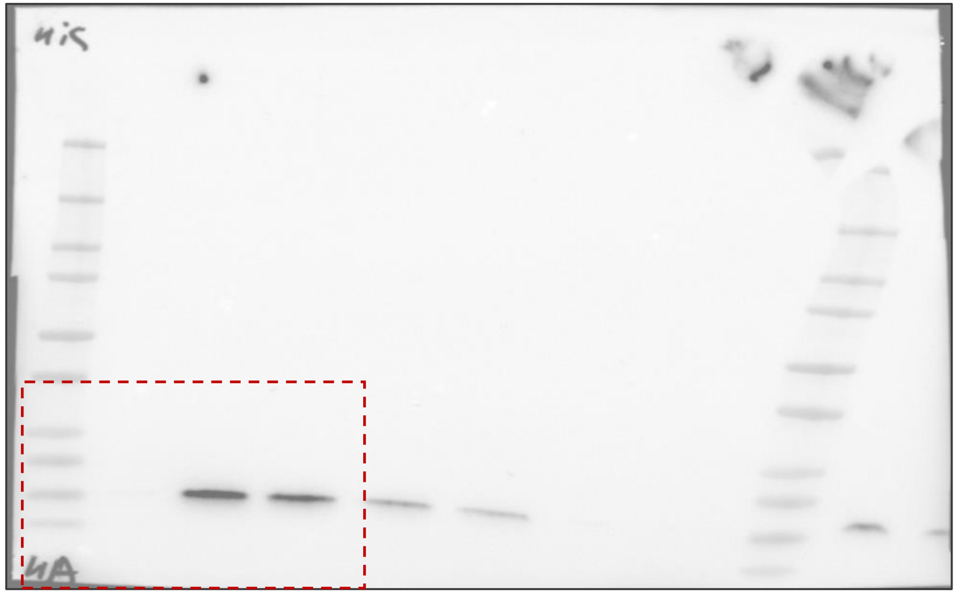
